# Verification of the phenylpropanoid pinoresinol biosynthetic pathway and its glycosides in *Phomopsis* sp. XP-8 using ^13^C stable isotope labeling and liquid chromatography coupled with time-of-flight mass spectrometry

**DOI:** 10.1101/354605

**Authors:** Yan Zhang, Junling Shi, Yongqing Ni, Yanlin Liu, Zhixia Zhao, Qianqian Zhao, Zhenhong Gao

## Abstract

*Phomopsis* sp. XP-8, an endophytic fungus from the bark of Tu-Chung (*Eucommiaulmoides*Oliv), revealed the pinoresinol diglucoside (PDG) biosynthetic pathway after precursor feeding measurements and genomic annotation. To verify the pathway more accurately, [^13^C_6_]-labeled glucose and [^13^C_6_]-labeled phenylalanine were separately fed to the strain as sole substrates and [^13^C_6_]-labeled products were detected by ultra-high performance liquid chromatography-quantitative time of flight mass spectrometry. As results, [^13^C_6_]-labeled phenylalanine was found as [^13^C_6_]-cinnamylic acid and *p*-coumaric acid, and [^13^C_12_]-labeled pinoresinol revealed that the pinoresinol benzene ring came from phenylalanine via the phenylpropane pathway. [^13^C_6_]-Labeled cinnamylic acid and *p*-coumaric acid, [^13^C_12_]-labeled pinoresinol, [^13^C_18_]-labeled pinoresinol monoglucoside (PMG), and [^13^C_18_]-labeled PDG products were found when [^13^C_6_]-labeled glucose was used, demonstrating that the benzene ring and glucoside of PDG originated from glucose. It was also determined that PMG was not the direct precursor of PDG in the biosynthetic pathway. The study verified the occurrence of the plant-like phenylalanine and lignan biosynthetic pathway in fungi.

**Importance:** Verify the phenylpropanoid-pinoresinol biosynthetic pathway and its glycosides in an endophytic fungi.

## Introduction

Pinoresinol diglucoside (PDG), (+)-1-pinoresinol 4, 4′ ‭di-O-β-D-glucopyranoside, is a major antihypertensive compound found in Tu-Chung, a traditional herb medicine with excellent efficacy for lowering blood pressure (1, 2). PDG possesses the potential to prevent osteoporosis (3). PDG is converted to enterolignans by intestinal microflora (4); thus, showing potential to reduce the risk of breast cancer (5) and other estrogen-dependent cancers (6).

PDG is found primarily in plants as lignans (1,7) but yields are very low. *Phomopsis* sp. XP-8 is an endophytic fungus isolated from the bark of Tu-Chung that was previously found to produce PDG *invitro* (8); thus, providing an alternative resource to obtain PDG. This is the first report on the capability of a microorganism to synthesize lignan. However, production was also very low. Therefore, it is essential to identify the PDG biosynthetic pathway in this strain.

The lignan biosynthetic pathway has only been reported in plants until now (9,10). Synthesis of pinoresinol (Pin) in plants occurs via oxidative coupling of monolignols, which are synthesized through the phenylpropanoid pathway with phenylalanine (Phe), cinnamicacid, *p*-coumarate, *p*-coumaroyl-CoA, caffeate, ferulate, feruloy-CoA, coniferylaldehyde, and coniferyl alcohol as intermediates or precursors (11, 12) (Fig.1). Pinoresinol monoglucoside (PMG) and PDG are converted from Pin by UDP-glucose-dependent glucosyltransferase (13). However, the biosynthesis of PDG from Pin has not been detected in plants and the Pin, PMG, and PDG biosynthetic pathways have not been illustrated in microorganisms.

**Fig.1.**
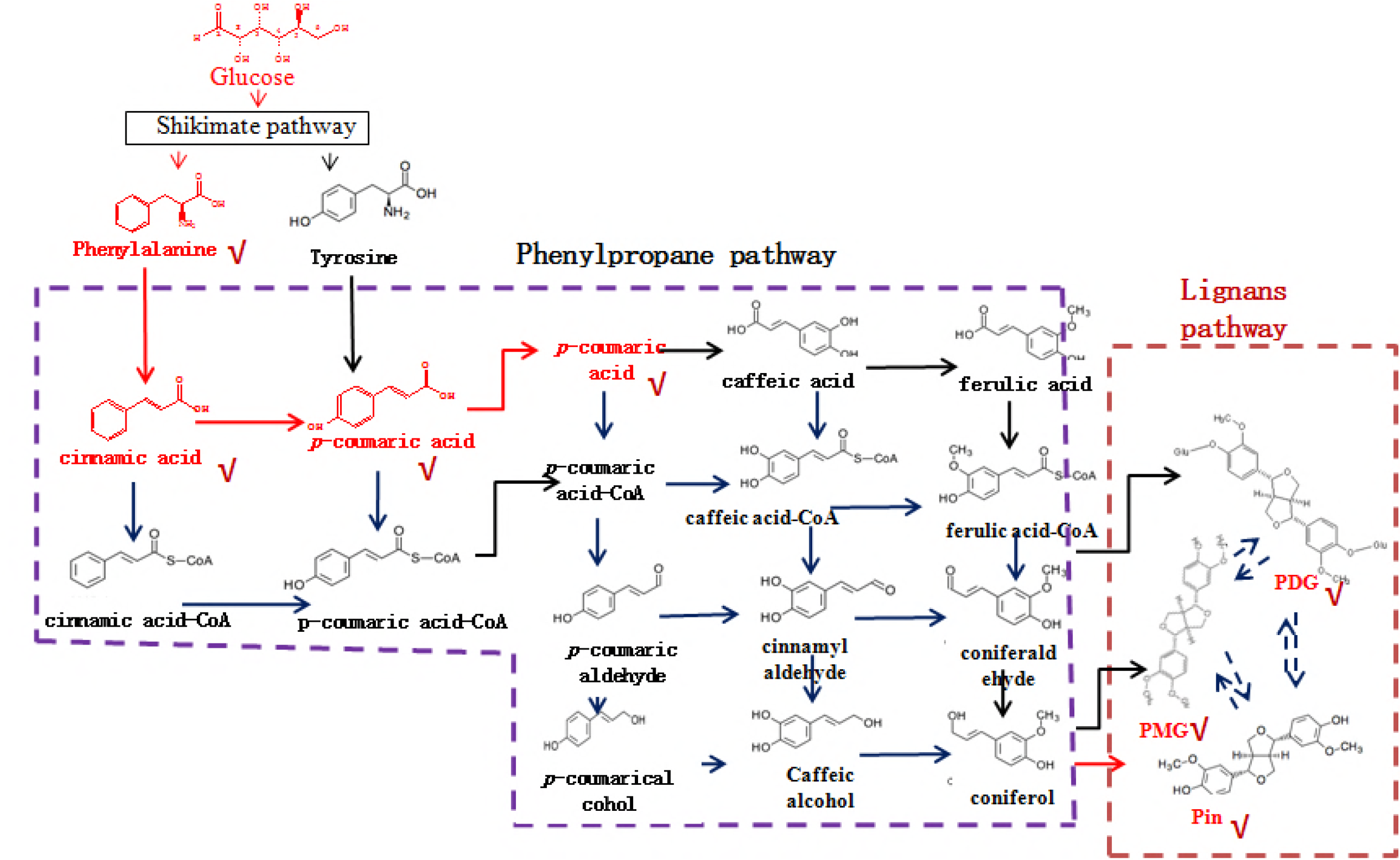
Biosynthetic pathways leading to lignans in plants.

We previously reported that *Phomopsis* sp. XP-8 converts mung bean starch and polysaccharides to Pin, PMG, and PDG. Phe, cinnamic acid, and *p*-coumaric acid have been detected as products of the bioconversion (14,15). Precursor feeding and enzymatic activity measurements indicate that this strain synthesizes PDG via many steps, such as during mass flow of the phenylpropanoid pathway (16). Genomic annotation indicates that the phenylpropane pathway exists in this strain (17) and some other microorganisms (18). However, the functions of the denoted genes have not been verified until now. Therefore, it is necessary to verify the entire PDG biosynthetic pathway in *Phomopsis* sp. XP-8.

Using stable or radioactive isotope-labeled compounds is an efficient and reliable strategy to verify the mass flow of unknown biosynthetic pathways by tracing the isotope-labeled compounds from substrates to products (19). ^13^C-labeled substrates have been used to shed light on the biodegradation pathways of organic pollutants (20). Isotope labeling combined with high-resolution mass spectrometry have also been used to track the abiotic transformation of pollutants in aqueous mixtures (21). In recent years, liquid chromatography-mass spectrometry (LC–MS) and ultra-high performance liquid chromatography (UPLC) systems have been developed to facilitate the analysis of many substances at the same time with high sensitivity and selectivity (22). Stable isotope-labeled compounds have also been employed in several areas of biomedical research (23). The combination of stable isotope-labeling techniques with MS has allowed rapid acquisition and interpretation of data and has been used in many fields, including distribution, metabolism, food, and excretion studies (24, 25, 26). The biochemical pathway of the aromatic compounds in tea has been also been revealed using the stable isotope labeling method (27).

In this study, we applied stable isotopes and MS to trace the PDG biosynthetic pathway. ^13^C_6_ stable isotope-labeled glucose and ^13^C_6_ Phe were used as the substrates and electrospray ionization-quantitative time of flight tandem mass spectrometry (ESI-Q-TOF-MS/MS) was used to identify the products.

## Materials and methods

### Microorganism and chemicals

*Phomopsis* sp. XP-8 previously isolated from the bark of Tu-Chung and stored at the China Center for Type Culture Collection (Wuhan, China) (code: *Phomopsis* sp. CCTCC M 209291) was used in the study.

Phe (purity ≥ 98%, Sigma, St. Louis, MO, USA), cinnamic acid and *p*-coumaric acid (purity ≥ 98%; Aladdin, Shanghai, China), PDG, PMG, and Pin (purity ≥ 99%; National Institutes for Food and Drug Control, Beijing, China) were used as the standards (dissolved in methanol) for the structural analysis and product identification.[^13^C_6_]-Labeled phenylalanine and glucose were purchased from the Qingdao TrachinoidCo (≥99%; Qingdao, China). The purity of the [^13^C_6_]-labeled Phe and glucose was 99%. Methanol (HPLC grade) was purchased from Fisher Scientific(Fairlawn, NJ, USA). The water used in the experiment was purified using a Milli-Q water purification system (18.5 M) (Millipore Corp., Bedford, MA, USA). Other reagents and chemicals were of analytical grade.

### Preparation of *Phomopsis* sp. XP-8 cells

*Phomopsis*sp.XP-8 was grown at 28°C on potato dextrose agar plates for 5 days. Then, three pieces of mycelia (5 mm in diameter) were inoculated into 100 mL liquid potato dextrose broth in a 90 250-mL flask and cultivated at 28°C on a rotary shaker (180 rpm). After 4 days, the cells were collected by centrifugation at 4°C (1,136×g for10 min) using a centrifuge (HC-3018R, Anhui USTC Zonkia Scientific Instruments Co., Ltd., Anhui, China). The cells were washed twice with sterile water and used for bioconversion according to the experimental design.

### Bioconversion systems

The bioconversion with normal glucose as the sole substrate was carried out in a 250-mL flask containing 100 mL of ultrapure water (pH 7),15 g/L glucose, and the prepared *Phomopsis*sp.XP-8 cell set a ratio of 10 g cells (wet weight) per 100 mL medium. To track the mass flow from glucose to PDG, 15 g/L glucose was changed to 5 g/L [^13^C_6_]-labeled glucose (5 g/L) in the above medium and the same conditions were used for bioconversion.

Bioconversion with Phe as the sole substrate was carried out in medium containing 0.15 g/L glucose (used for the glycosyl donors), 7 mM [^13^C_6_]-labeled phenylalanine, and the prepared *Phomopsis* sp. XP-8 cells at a ratio of 10 g wet cells per 100 mL medium.

All bioconversions were carried out for 48 h at 28°C and 180 rpm. At the end of bioconversion, the broth was collected and filtered through an intermediate speed qualitative filter paper before the products were detected.

### Identification of the accumulated products during bioconversion

The products were extracted from the vacuum-evaporated (0.09 MPa, 50°C) bioconversion broth with methanol and adjusted to 4 mL for the UPLC measurements after filtration through a membrane (0.45 μm, 13 mm diameter; Millipore, Billerica, MA, USA).The UPLC analysis was performed on a Waters Acquity UPLC system (Waters Corp., Milford, MA, USA), equipped with a binary pump, a thermostatically controlled column compartment, and a UV detector. Gradient elution was performed on an Acquity UPLCTM BEH C18 column (50 mm × 2.1 mm I.D., 1.7 m; Waters) and the column temperature was maintained at 30°C, while sample temperature was 10°C (15).

The MS analysis of the products was carried out on a Q-TOF Premier^™^ with an ESI source(Waters Corp.) at the optimized parameters of: capillary voltage, 2.8 kV; sampling cone voltage, 20 V; extractor voltage, 4 V; source temperature, 100°C; desolvation temperature, 250°C, and flow rate of the desolvation gas (N2), 400 L/h. The collision cell parameters for the Q-TOF-MS/MS analysis were: collision gas (Argon) flow rate, 0.45 L/h; collision energy, 15–35 eV. The mass spectra were recorded using full scan mode over a mass range of m/z 100–800in negative ion mode. The MS acquisition rate was set to 1.0 s, with a 0.02 s inter-scan delay. The Q-TOF-MS/MS experiments were carried out by setting the quadrupole to allow ions of interest to pass prior to fragmentation in the collision cell.

Accurate mass measurements were obtained by means of a lock mass that introduces a low flow rate (3 L/min) of a chrysophanol (253.0499) calibrating solution in the ESI-Q-TOF-MS and ESI-Q-TOF-MS/MS. All operations and acquisition and data analyses were controlled by Masslynx V4.1 software (Waters Corp.).

### Data processing

Peak detection, alignment, and identification of the detected compounds were performed using Masslynx V4.1 software (Waters Corp.).The MS/MS fragmentation patterns were used for informative non-targeted metabolic profiling of the LC-MS data, and the acquired LC-MS/MS spectrum was identified after comparison with spectra proposed by the Mass bank database (www.massbank.jp), the KEGG database, and related reports.

## Results

### Detection of products converted from normal glucose

Production of PDG, monoglucoside (PMG), Pin, Phe, *p*-coumaric acid (*p*-Co), and cinnamic acid (Ca) were detected in bioconversion systems using glucose as the sole substrate. Data in Figs. 2–7 show the mass spectra of these compounds accumulated in the bioconversion systems and the corresponding standards.

**Fig.2.**
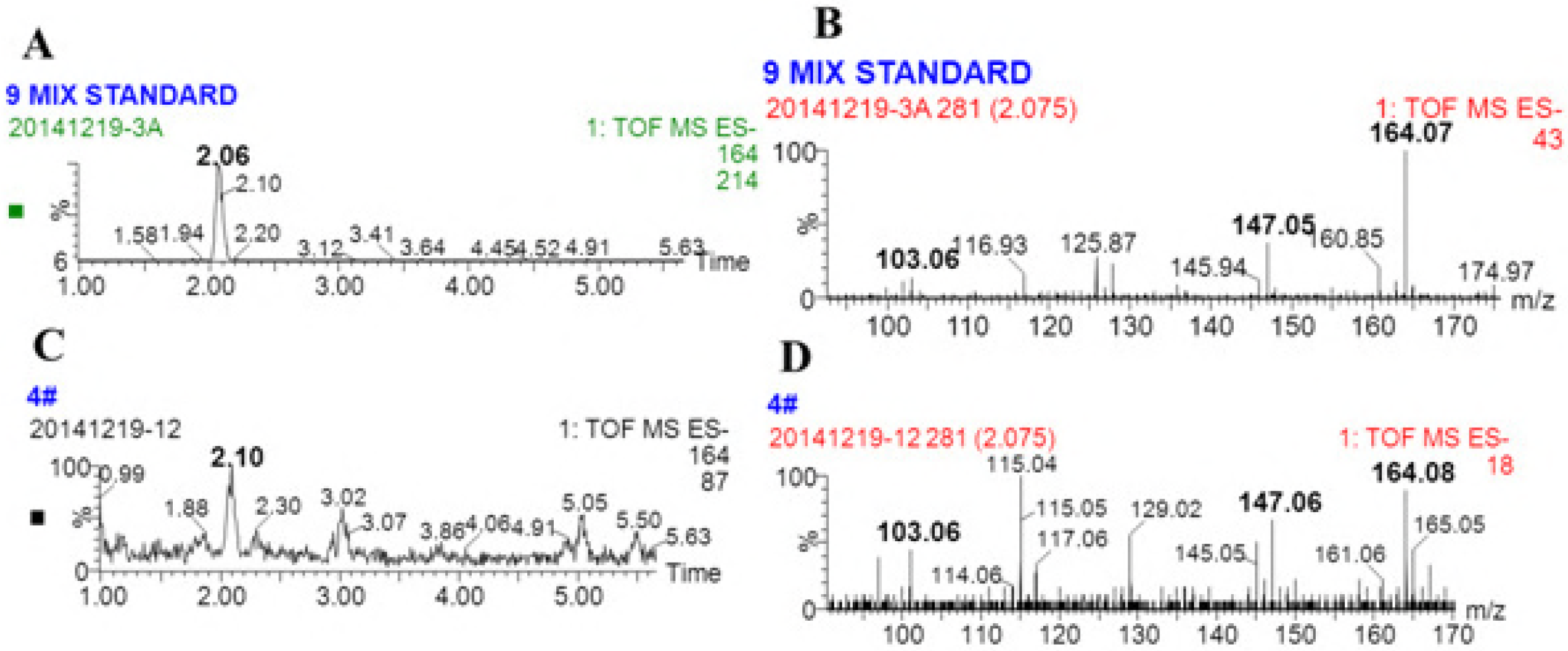
Total ion current chromatogram and mass spectrum of standard phenylalanine and that in the samples. (A and B show the total ion current chromatogram and the mass spectrum of standard phenylalanine, respectively; C and D show the total ion current chromatogram and the mass spectrum of phenylalanine in the samples, respectively. Ion reaction was set tom/z=164)

**Fig.3.**
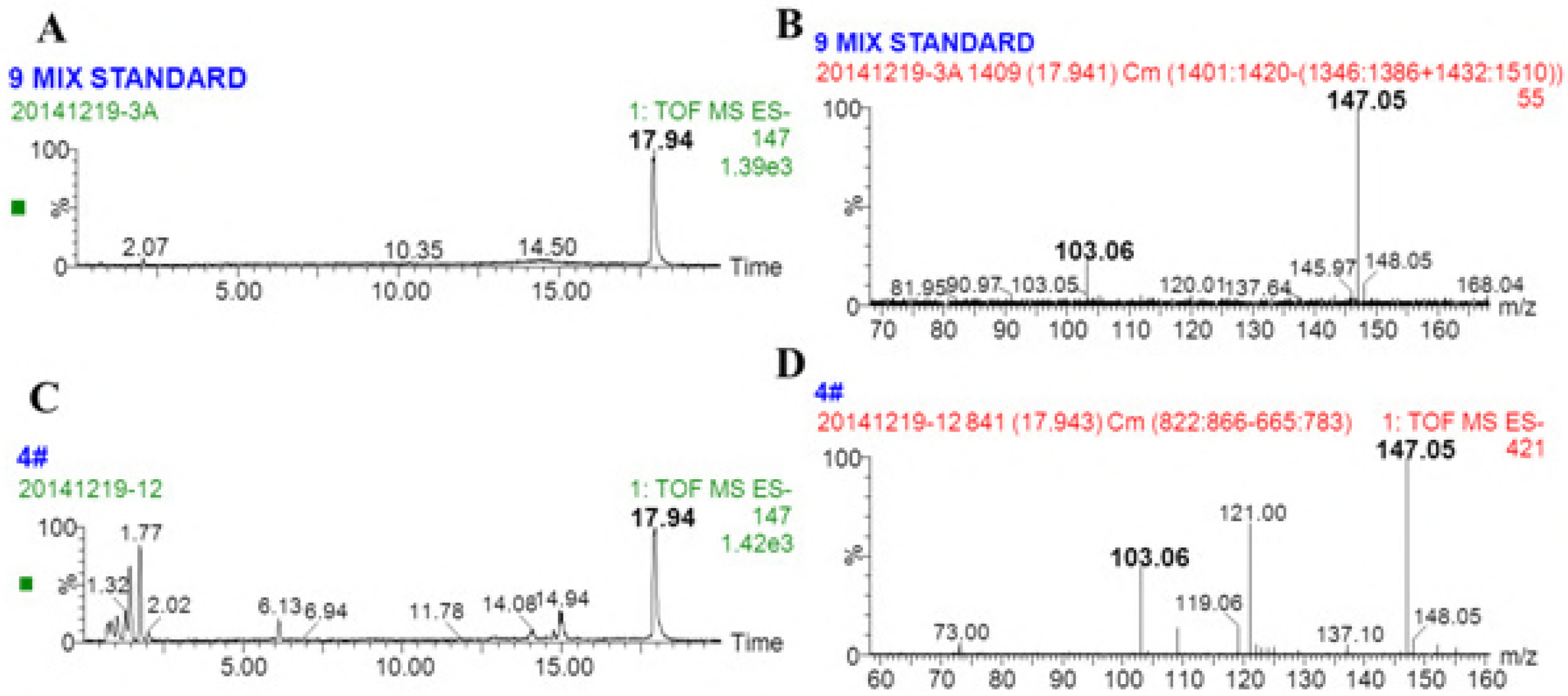
Total ion current chromatogram and mass spectrum of standard cinnamic acid and that in the samples. (A and B show the total ion current chromatogram and the mass spectrum of standard cinnamic acid, respectively; C and D show the total ion current chromatogram and the mass spectrum of the samples, respectively. Ion reaction was set to m/z=147)

**Fig.4.**
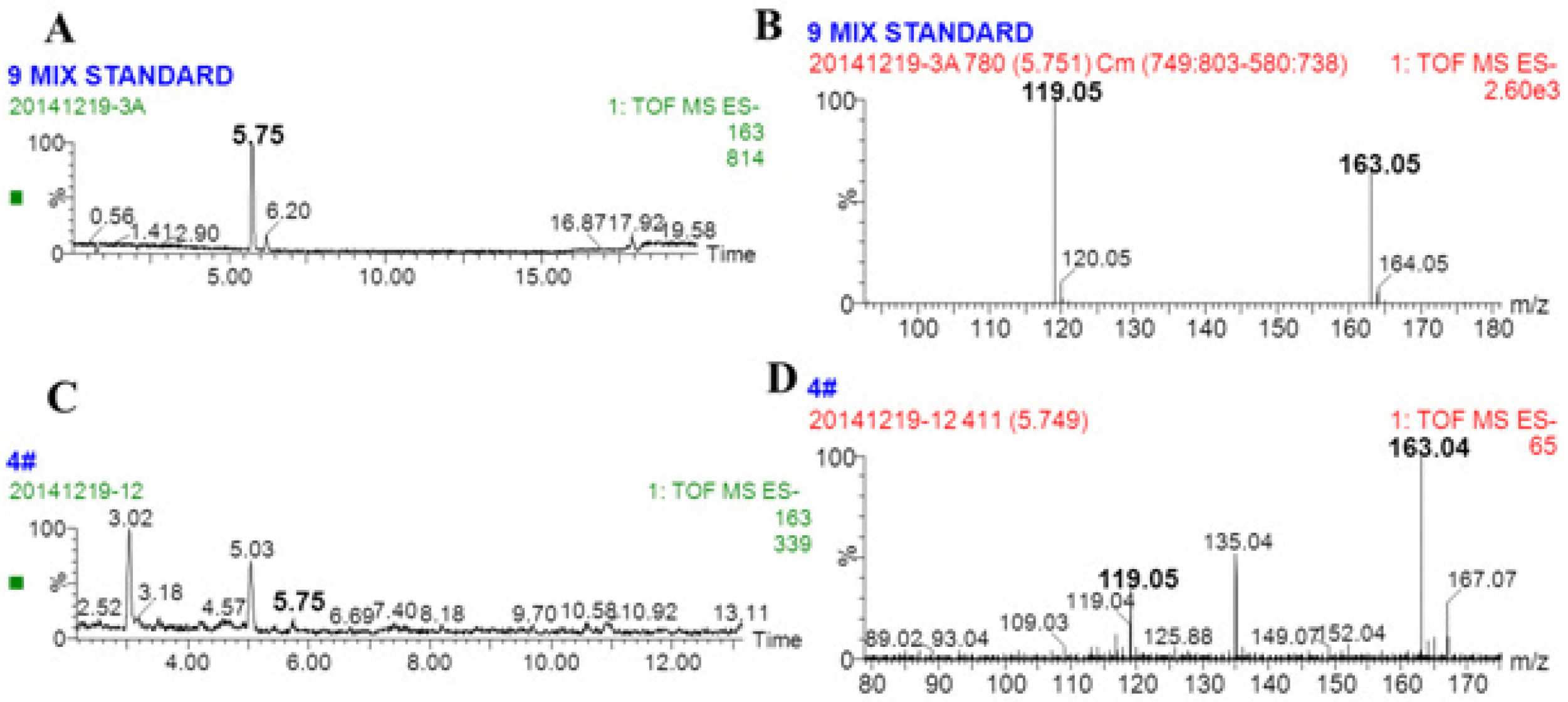
Total ion current chromatogram and mass spectrum of standard *p*-coumaric acid and that in the samples. (A and B show the total ion current chromatogram and the mass spectrum of standard *p*-coumaric acid, respectively; D-F show the total ion current chromatogram, and mass spectrum of *p*-coumaric acid in the samples, respectively. Ion reaction was set tom/z=163)

**Fig.5.**
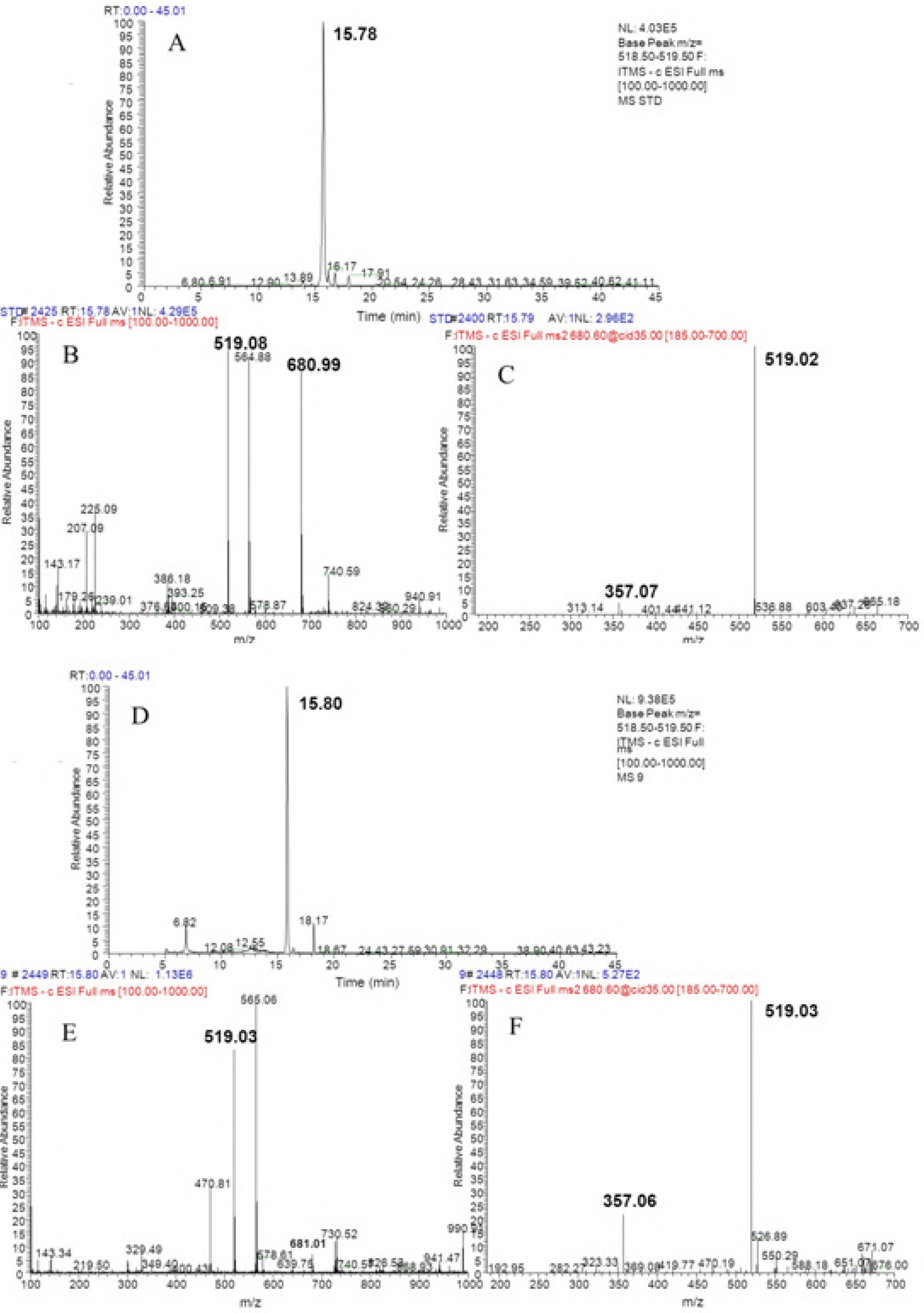
Total ion current chromatogram and mass spectrum of standard PDG and that extracted from samples. (A-C are the total ion current chromatogram, precursor ions, and daughter ions of standard PDG, respectively; D-F are the total ion current chromatogram, precursor ions, and daughter ions of the samples, respectively).

**Fig.6.**
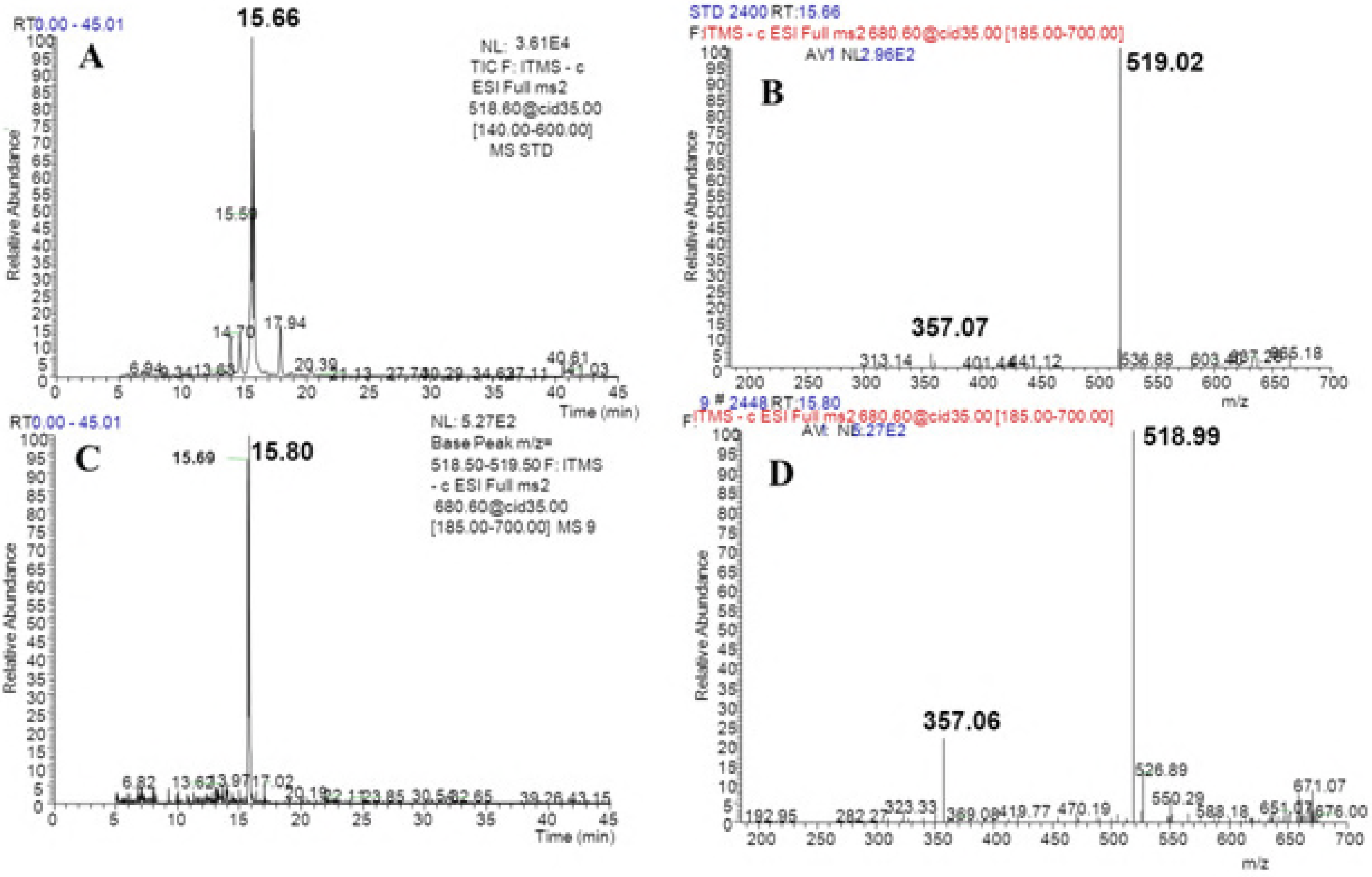
Total ion current chromatogram and mass spectrum of standard pinoresinol-4-O-β-Dglucopyranosideand and that in the samples. (A and B show the total ion current chromatogram and mass spectrum of standardpinoresinol-4-O-β-D-glucopyranoside, respectively; D-F show the total ion current chromatogram and the mass spectrum ofpinoresinol-4-O-β-D-glucopyranosideinsamples, respectively. Ion reaction was set tom/z=518.5-519.5)

**Fig.7.**
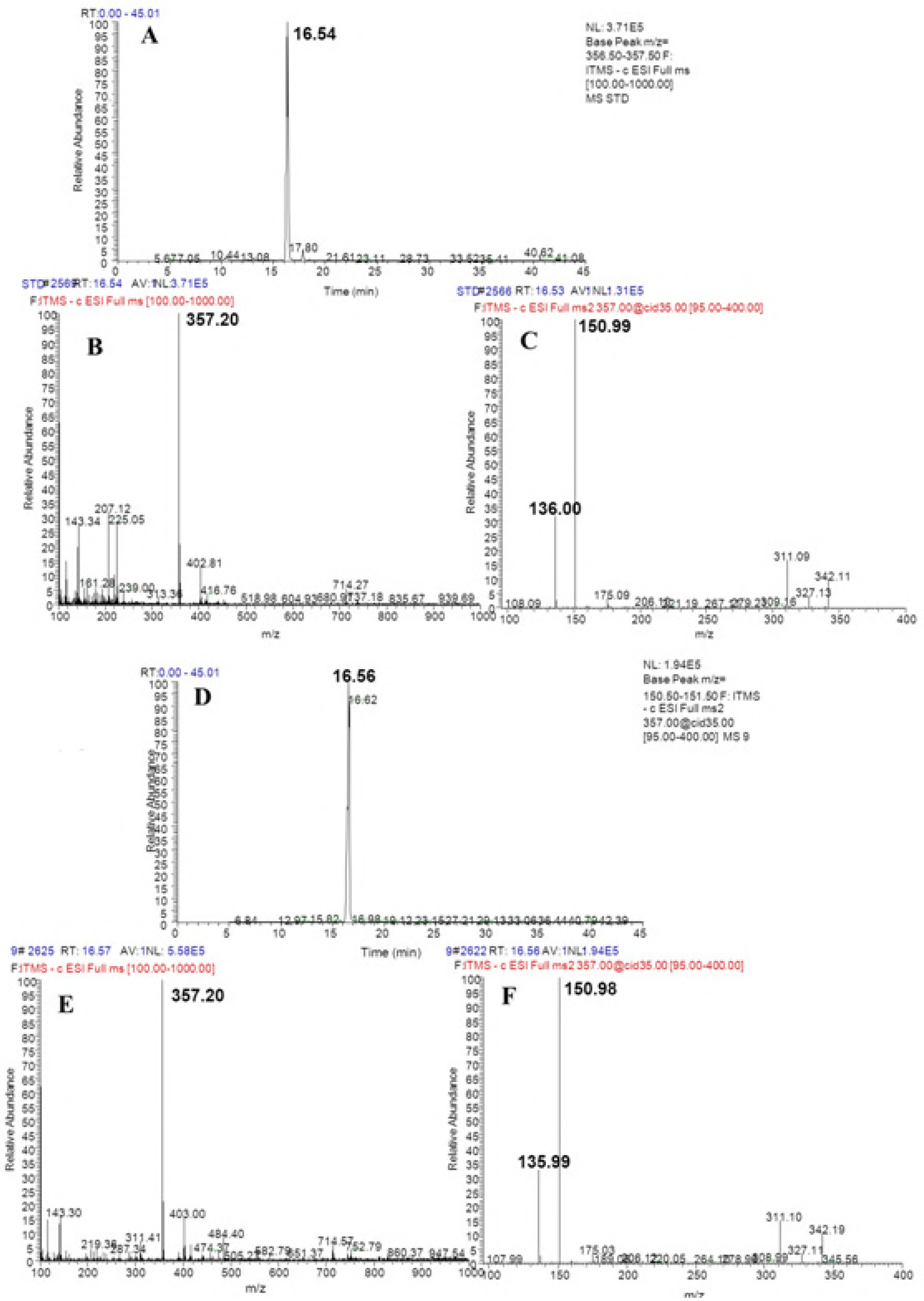
Total ion current chromatogram and mass spectrum of standard pinoresinol and that in the samples. (A-C show the total ion current chromatogram, precursor ions, and daughter ions of the pinoresinol standard, respectively; D-F show the total ion current chromatogram, precursor ions, and daughter ions of pinoresinol in the samples, respectively. Ion reaction was set tom/z=356.5-357.5)

Production of Phe was detected at a molecular weight and major daughter ions of m/z = 164.08 and m/z = 147.06, respectively (Fig. 2D),which was consistent with the data obtained from the corresponding standards (Fig. 2B). Similarly, production of PDG, PMG, Pin, *p*-Co, and CA was also detected in the bioconversion system, indicating that glucose was converted to these products by *Phomopsis* sp. XP-8, as only glucose was provided in the bioconversion system.

### Identification of products converted from[^13^C_6_]-labeled phenylalanine

The phenylpropane pathway in plants starts with Phe and ends with *p*-Co. The same mass flow was previously detected during production of PDG from glucose by *Phomopsis*sp.XP-8 (Zhang et al., 2015b). To verify this finding and the role of the Phe pathway in the biosynthesis of PDG, PMG, and Pin,[^13^C_6_]-labeled Phe was used as the sole substrate in the bioconversion system with 5 g/L glucose (mainly used as the glucoside donor). As results, ^13^C labeled Pin, Phe, *p*-Co, and Ca were successfully detected (Fig. 8).

**Fig.8.**
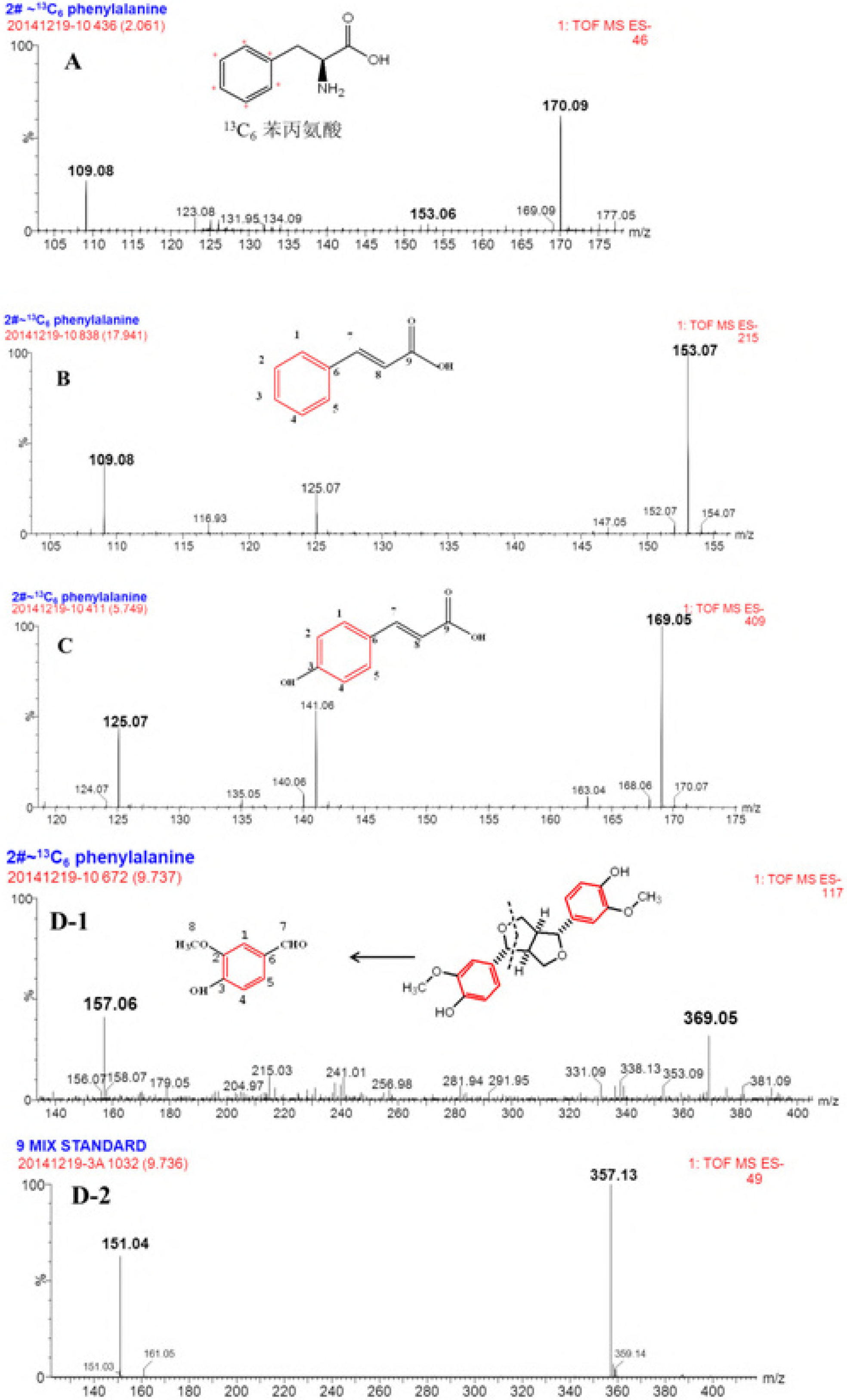
Mass spectrum of phenylalanine, cinnamic acid, *p*-coumaric acid, PDG, PMG, and Pin in the resting cell system using phenylalanine with the ^13^C_6_ stable isotope labeled as the substrate. (A: phenylalanine; B: cinnamic acid; C: *p*-coumaric acid; D: Pin)

The retention time (RT) of [^13^C_6_]-labeled Phe was the same as the Phe standard (Figs.2A and 8A). The molecular weight and major daughter ions of [^13^C_6_]-labeled Phe were obtained at m/z = 170.09 (Fig. 8A), indicating six ^13^C in Phe. Similarly, the other products were also successfully detected at the same RT of their corresponding normal standard substrates. All ^13^C-labeled product data and their corresponding standard substrates are summarized in Table 1.

**Table 1.**
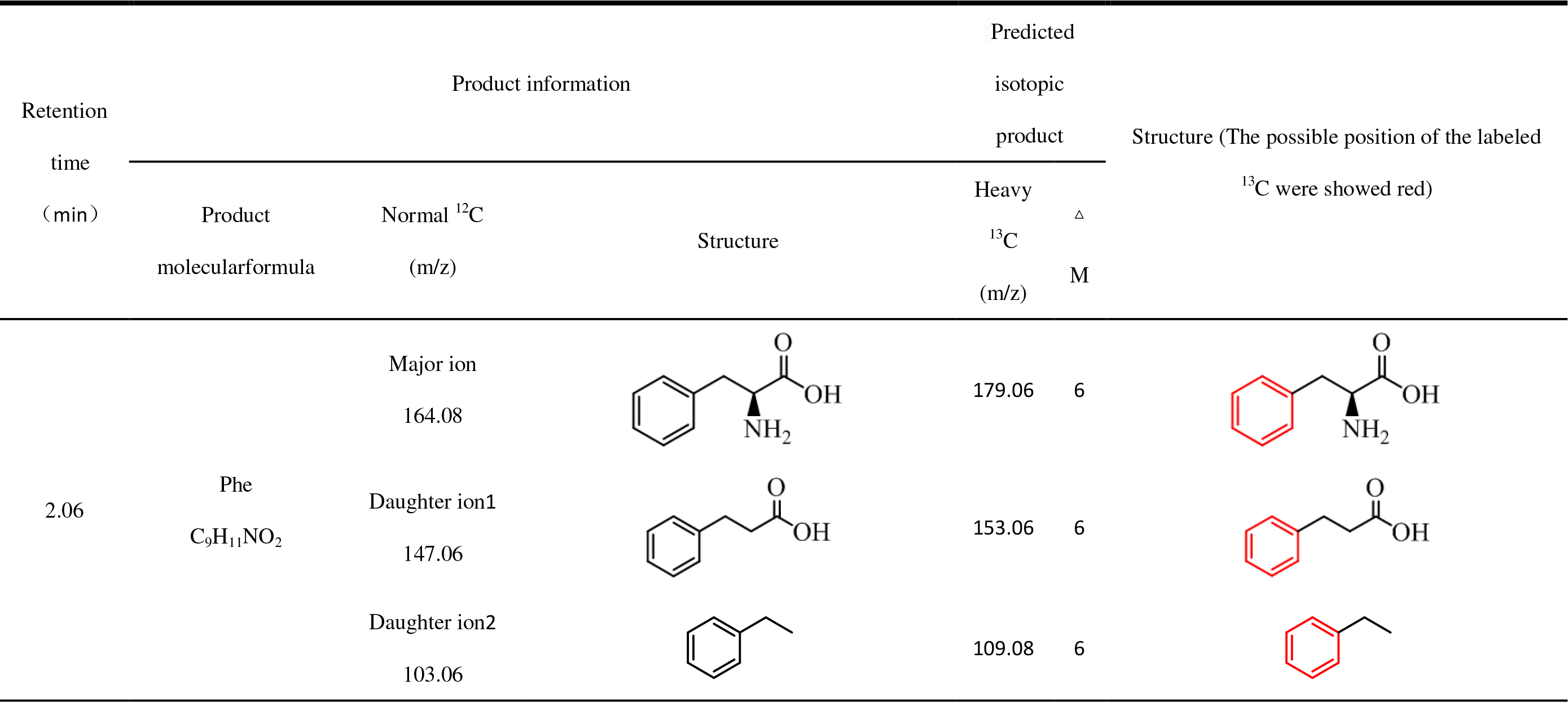
Predicted products with [^13^C_6_]-labeled phenylalanine as the substrate.

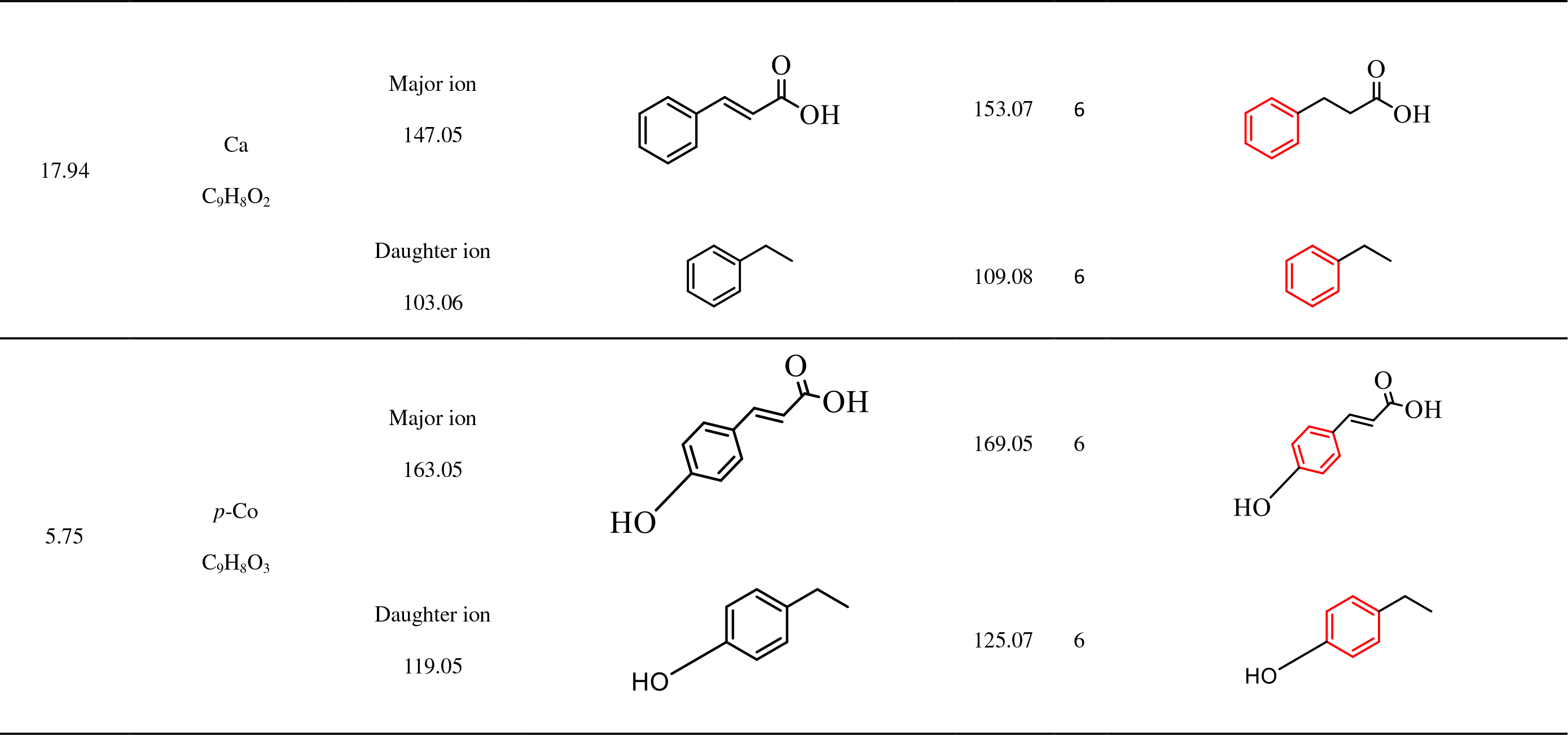

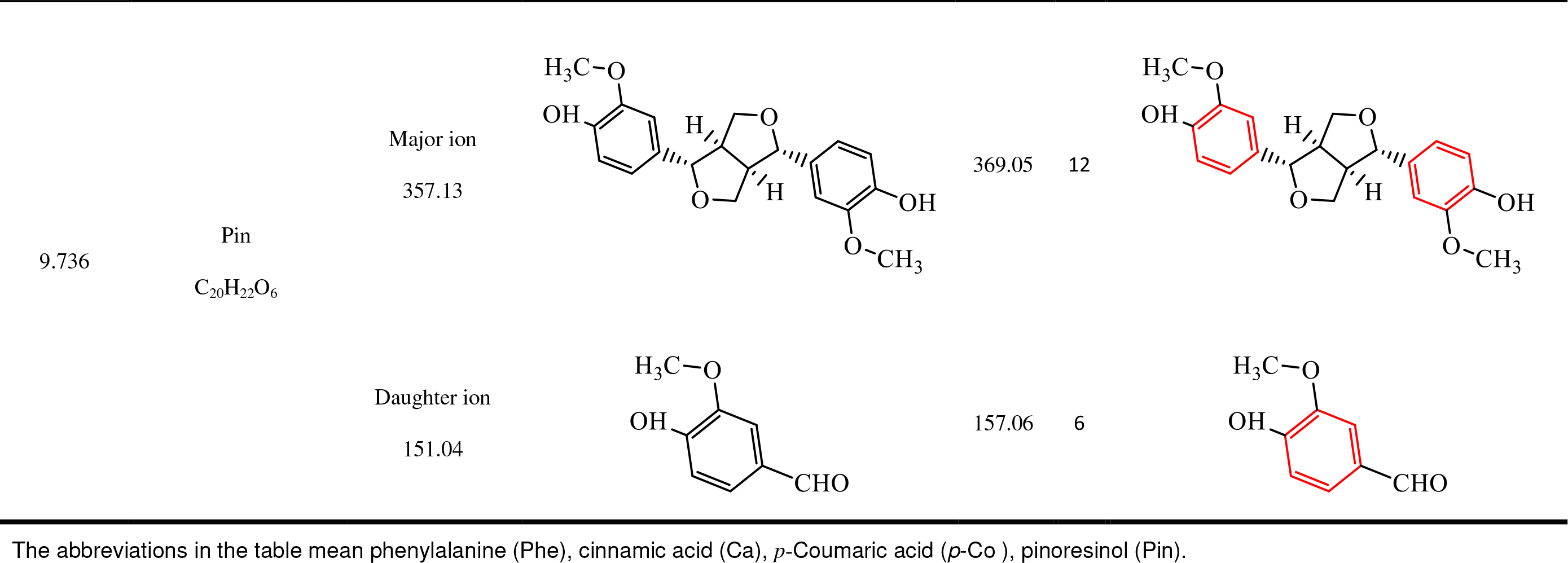

Standard Ca (C_9_H_8_O_2_) was detected at a RT of 17.94 min with molecular weight and major daughter ions obtained at m/z = 147.05 and m/z = 103.06, respectively. (Fig. 3A,B). ^13^C-Labeled Ca was detected in the bioconversion system at a molecular weight of m/z = 153.07, indicating that six ^13^C were introduced into Ca (Fig. 8B). The major daughter ions of ^13^C-labeled Ca were obtained at m/z 160 = 109.08, indicating six ^13^C referring to the standard Ca (m/z=103.06). The structure of ^13^C-labeled Ca with out-COOH was the major daughter ion at m/z=103.06. Therefore, it was deduced that the six ^13^C were in a benzene ring not in -COOH.

According to the mass spectra of the *p*-Co standard (C_9_H_8_O_3_, RT= 5.75min, the molecular weight and major daughter ions were obtained at m/z = 163.05 and m/z = 119.06 respectively) (Fig. 4A, B). *p*-Co produced in the conversion system was detected at a molecular weight of m/z= 169.05 and revealed six ^13^C by consulting the *p*-Co standard (m/z= 163.05). The major daughter ions of ^13^C-labeled *p*-Co were obtained at m/z =125.07,which was six more than that of the *p*-Co standard (m/z=119.06). The structure of ^13^C-labeled *p*-Co without-COOH was detected at the major daughter ion of m/z=119.06. Therefore, it was deduced that the six ^13^C might be distributed in the benzene ring.

^13^C-Labeled Pin was detected (Fig. 8D-1) and compared with the mass spectra of the Pin standard (C_20_H_22_O_6_, RT= 9.736 min, the molecular weight and major daughter ions were obtained at m/z = 357.13and m/z = 151.04 respectively) (Fig. 8D-2). The molecular weight of ^13^C-labeled Pin was identified at m/z= 369.05, which was 12 more than that of standard Pin (m/z= 357.13). The major daughter ions of ^13^C-labeled Pin were obtained at m/z =125.07, indicating six more than that of the Pin standard (m/z=151.04). The structure of ^13^C-labeled Pin with loss of a benzene ring was identified as the major daughter ion of m/z=151.04 (Fig. 7D-1). This result confirmed that the six^13^C were distributed in a benzene ring, whereas the other six ^13^C might be in a symmetrical benzene ring. Therefore, we deduced that the Pin with 12 ^13^C was bio-converted from the[^13^C_6_]-labeled Phe, Ca, or/and *p*-Co. This finding also confirmed that the benzene ring in Pin came from Phe, which is consistent with that of the lignan biosynthetic pathway in plants.

### Identification of products converted from [^13^C_6_]-labeled glucose

To explore where Phe originated from the Pin biosynthetic pathway, [^13^C_6_]-labeled glucose was supplied as the sole substrate in the bioconversion system with *Phomopsis* sp. XP-8 cells. As results, ^13^C labeled PDG, PMG, Pin, Phe, *p*-Co, and Ca were detected (Fig. 8).The products were detected according to the RTs of the corresponding standards. The detailed information on the products and corresponding standards is shown in Table 2.

**Table 2.**
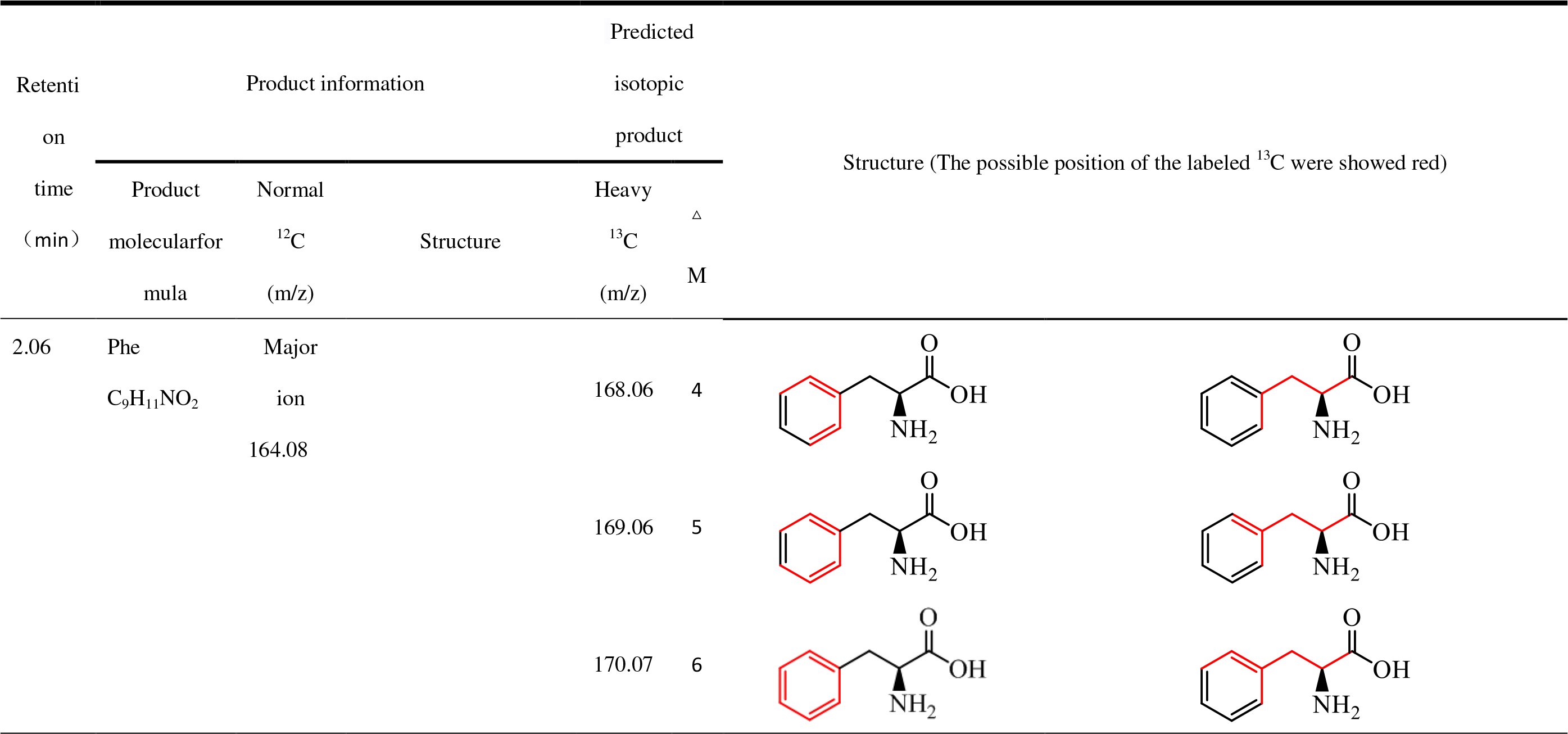
Predicted products with [^13^C_6_]-labeled glucose as the substrate.

| Retenti<br>on<br>time<br>( min ) | Product information |  |  | Predicted isotopic product |  | Structure (The possible position of the labeled <sup>13</sup>C were showed red) |
| --- | --- | --- | --- | --- | --- | --- |
|  | Product molecular formula | Normal <sup>12</sup>C (m/z) | Structure | Heavy <sup>13</sup>C (m/z) | Δ M |  |
| 2.06 | Phe<br>C<sub>9</sub>H<sub>11</sub>NO<sub>2</sub> | Major ion<br>164.08 |  | 168.06 | 4 |  |
|  |  |  |  | 169.06 | 5 |  |
|  |  |  |  | 170.07 | 6 |  |

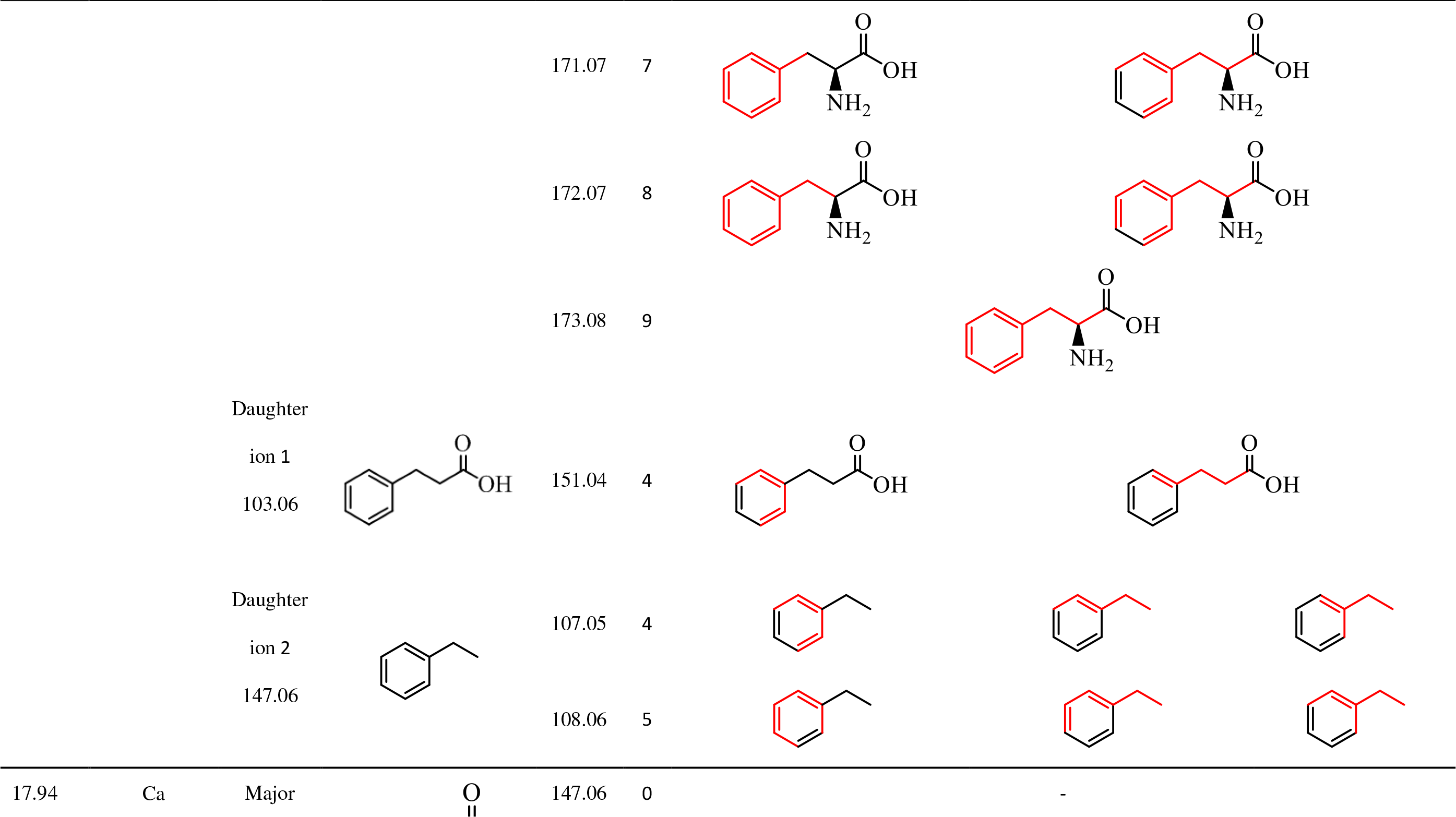

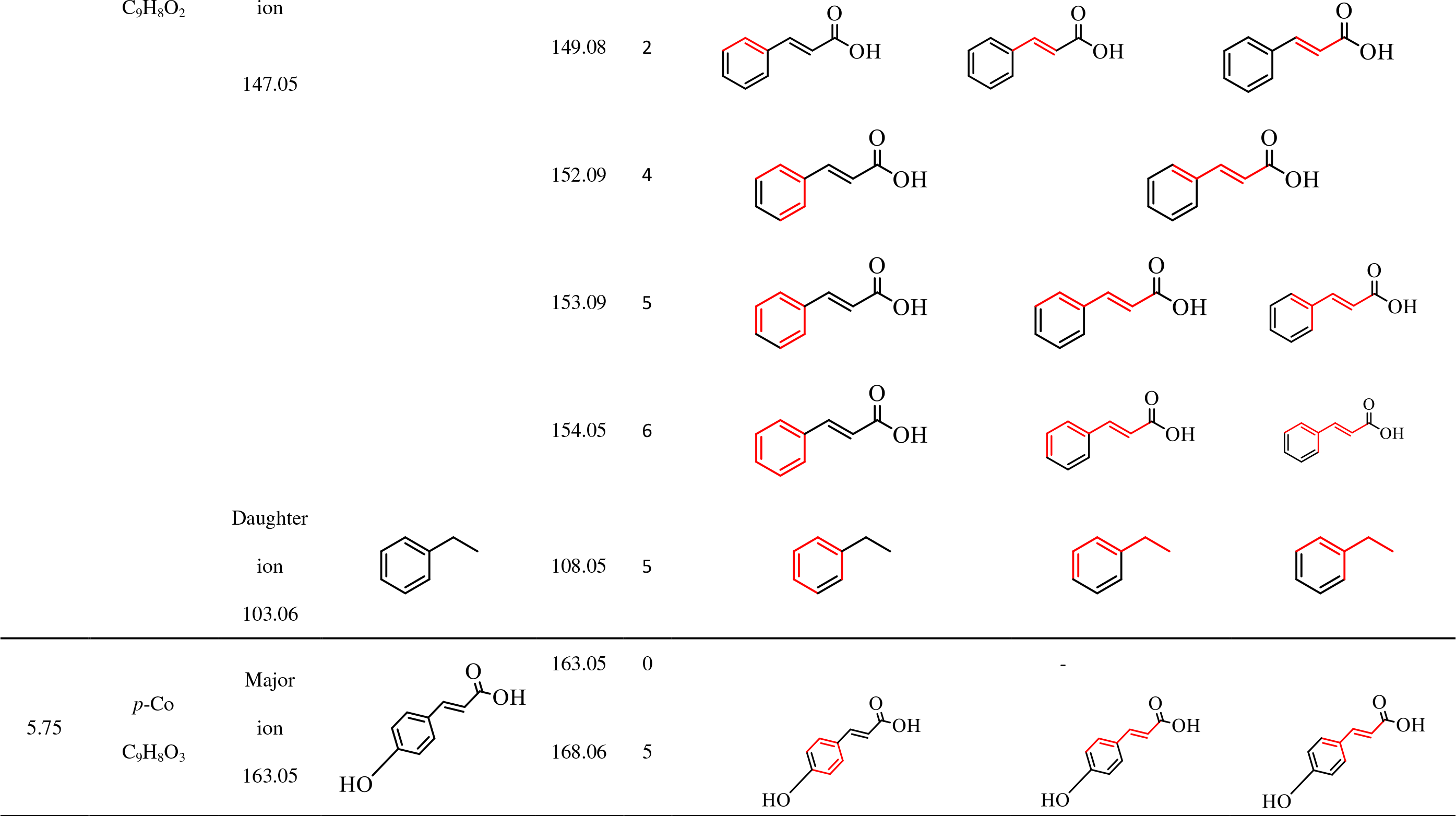

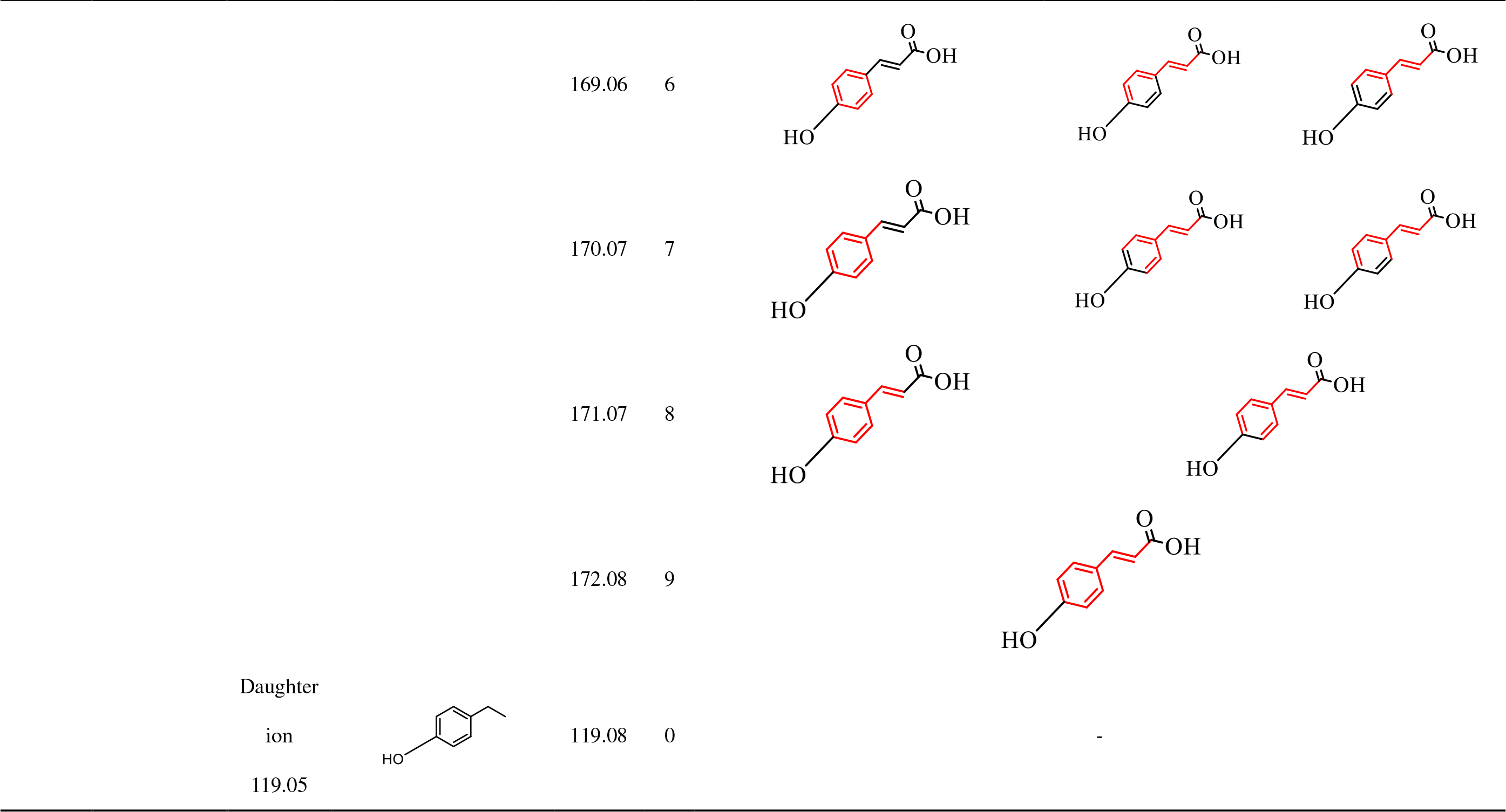

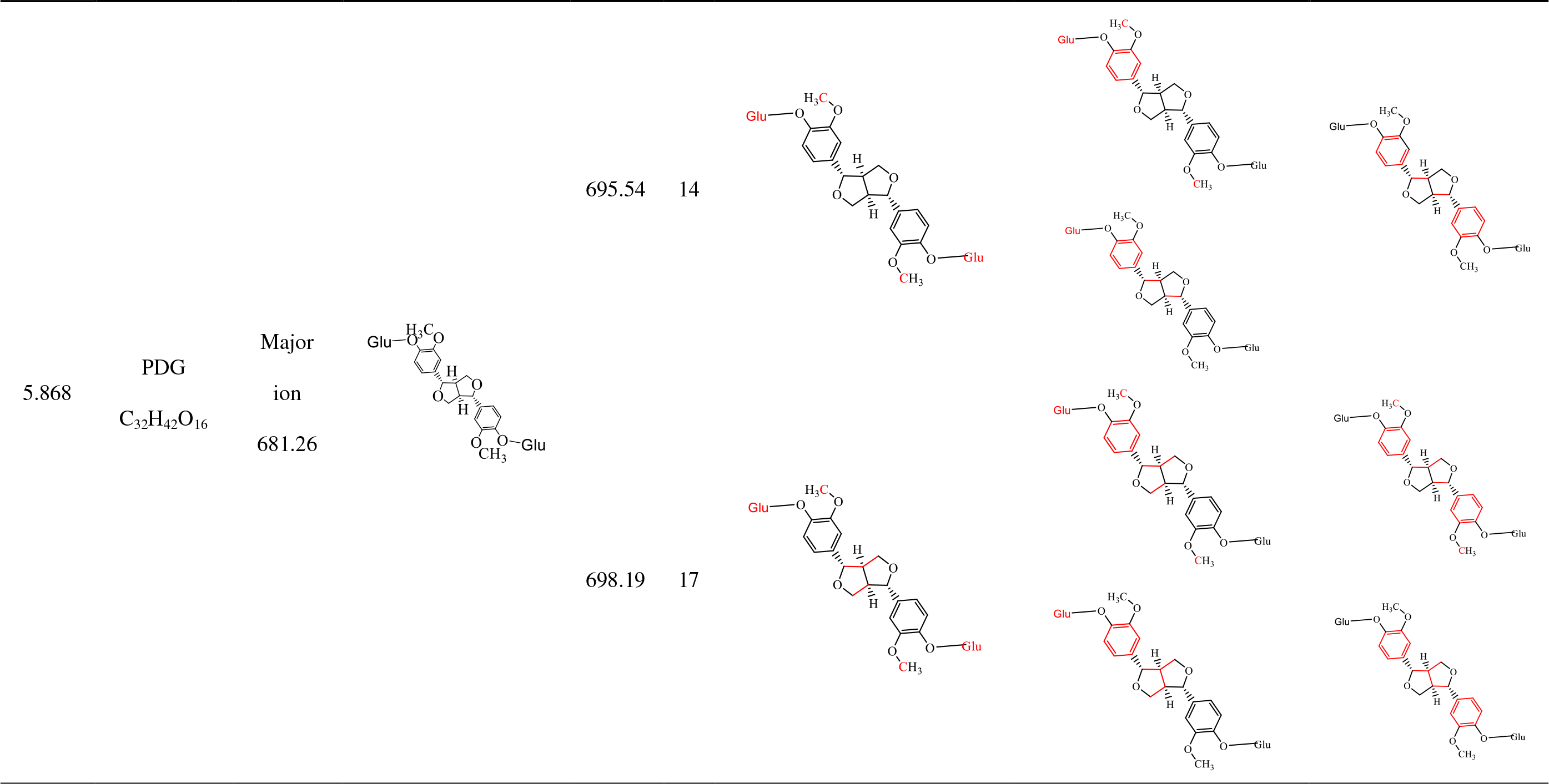

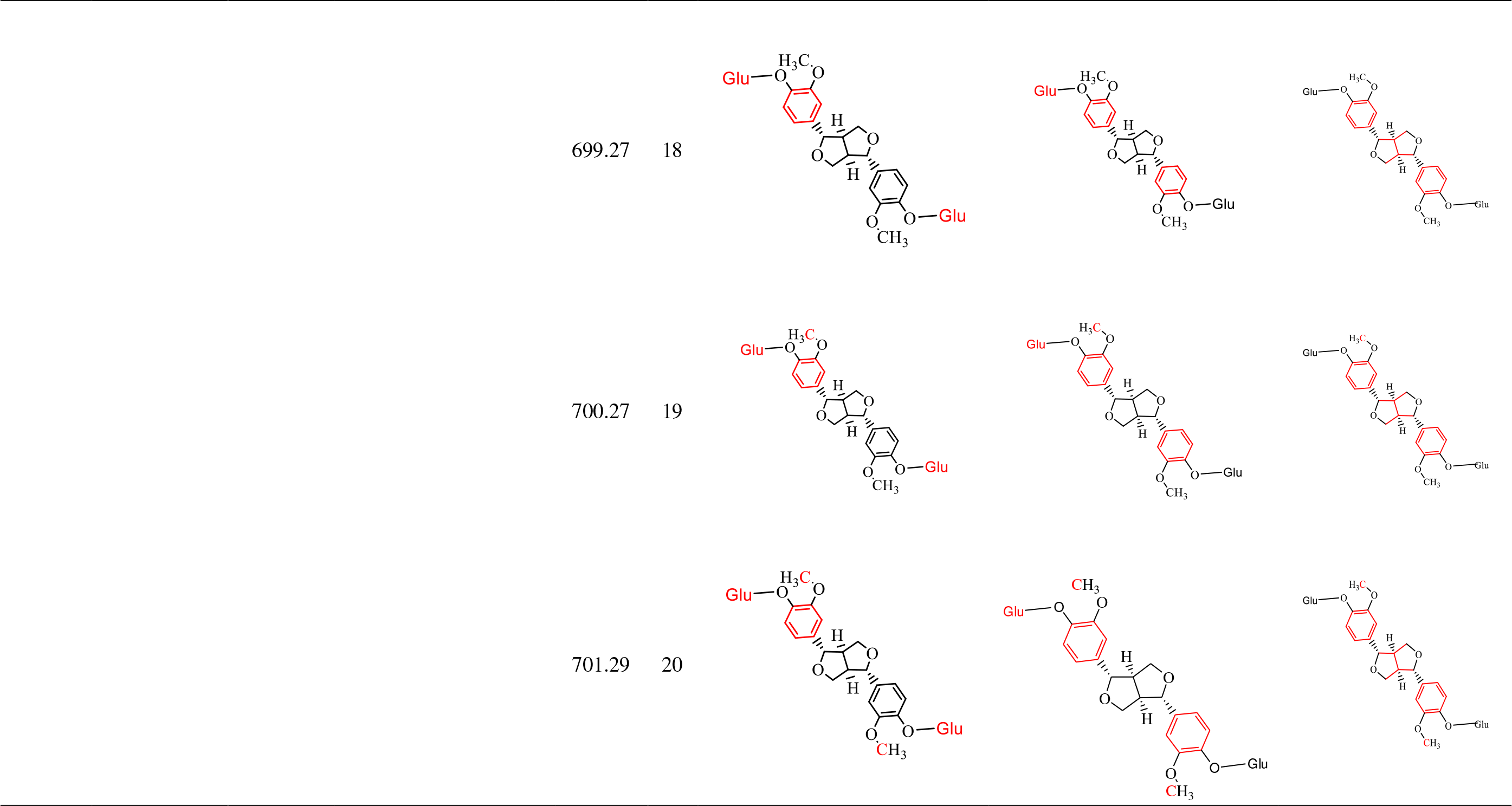

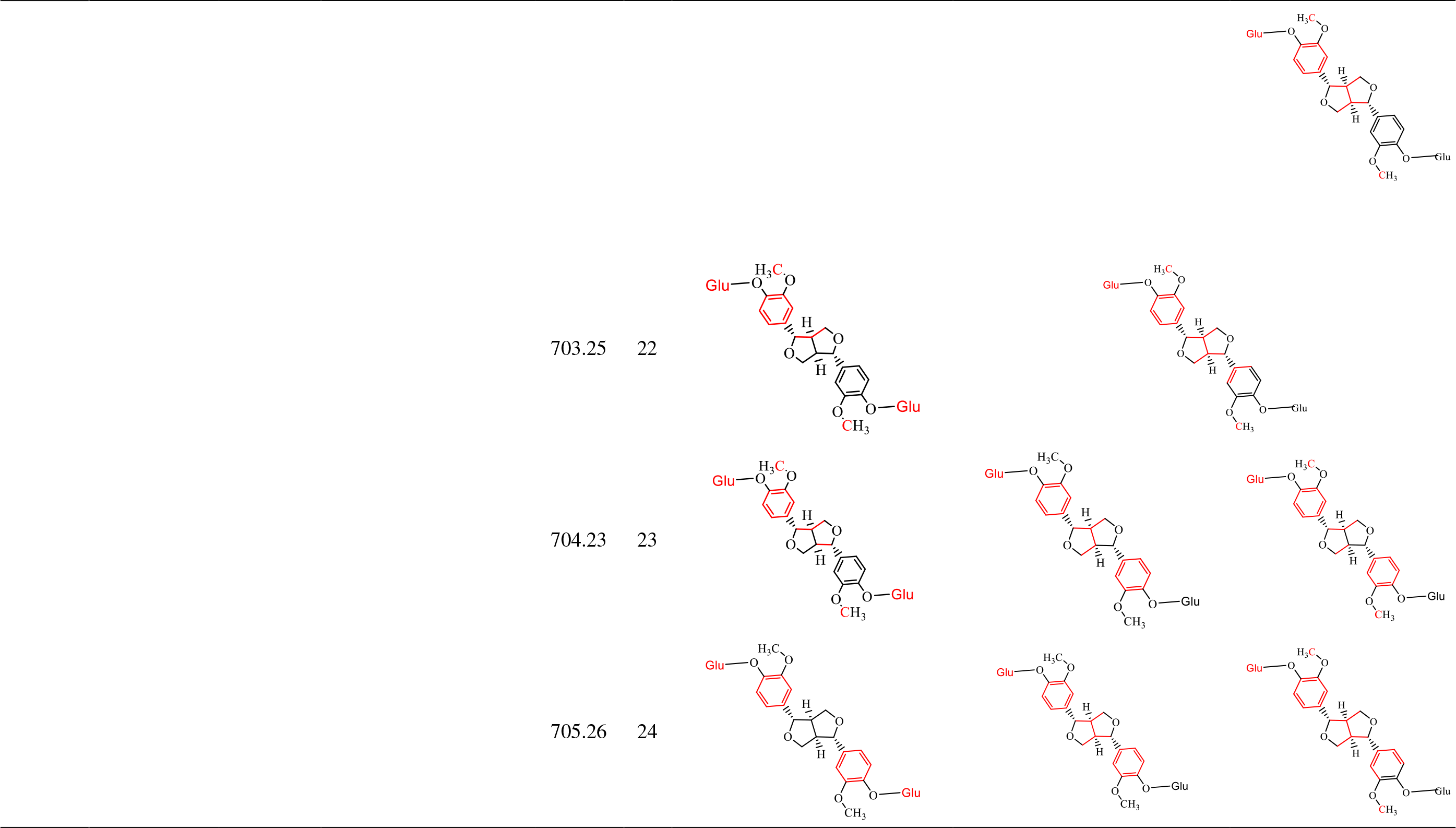

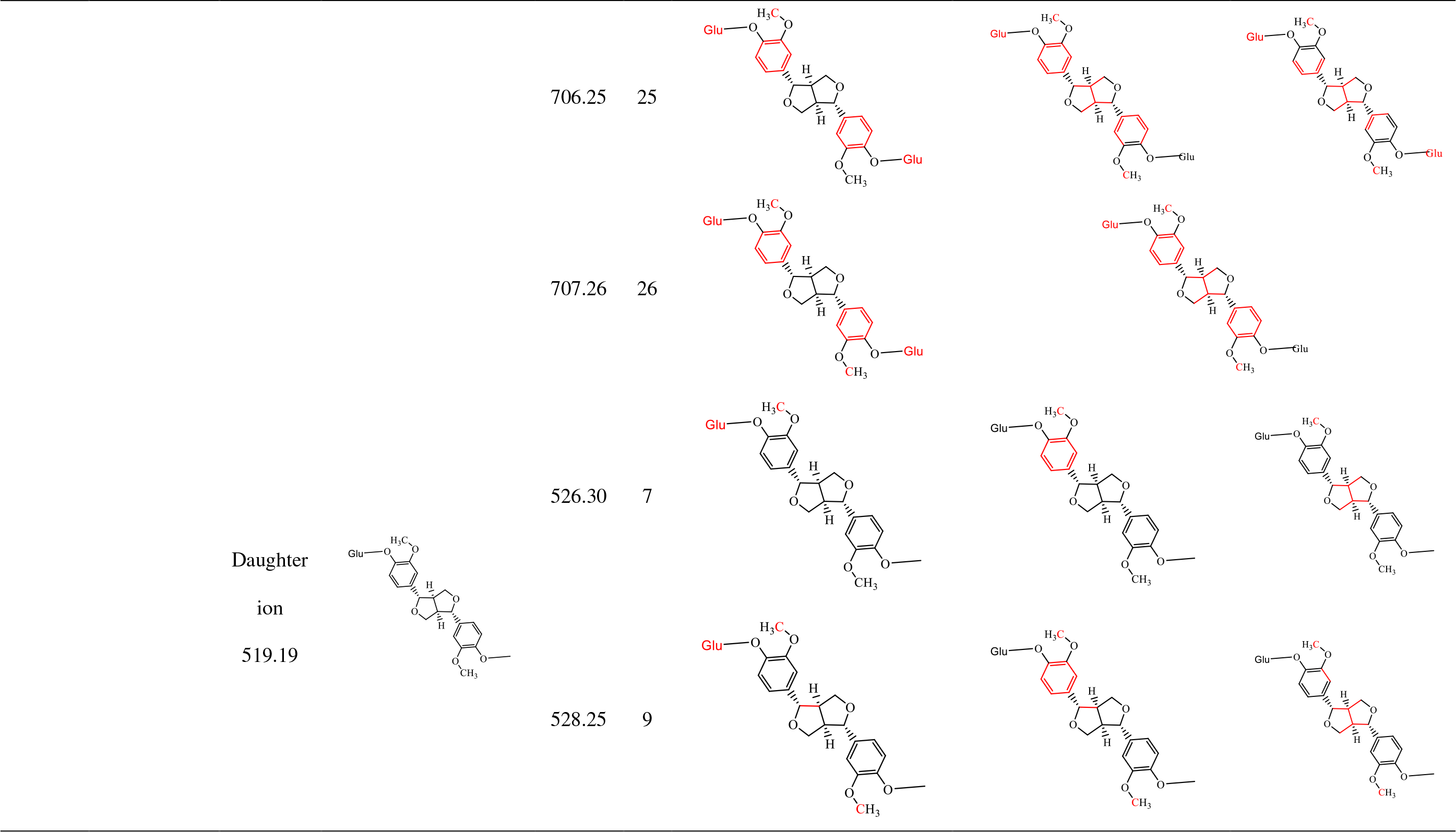

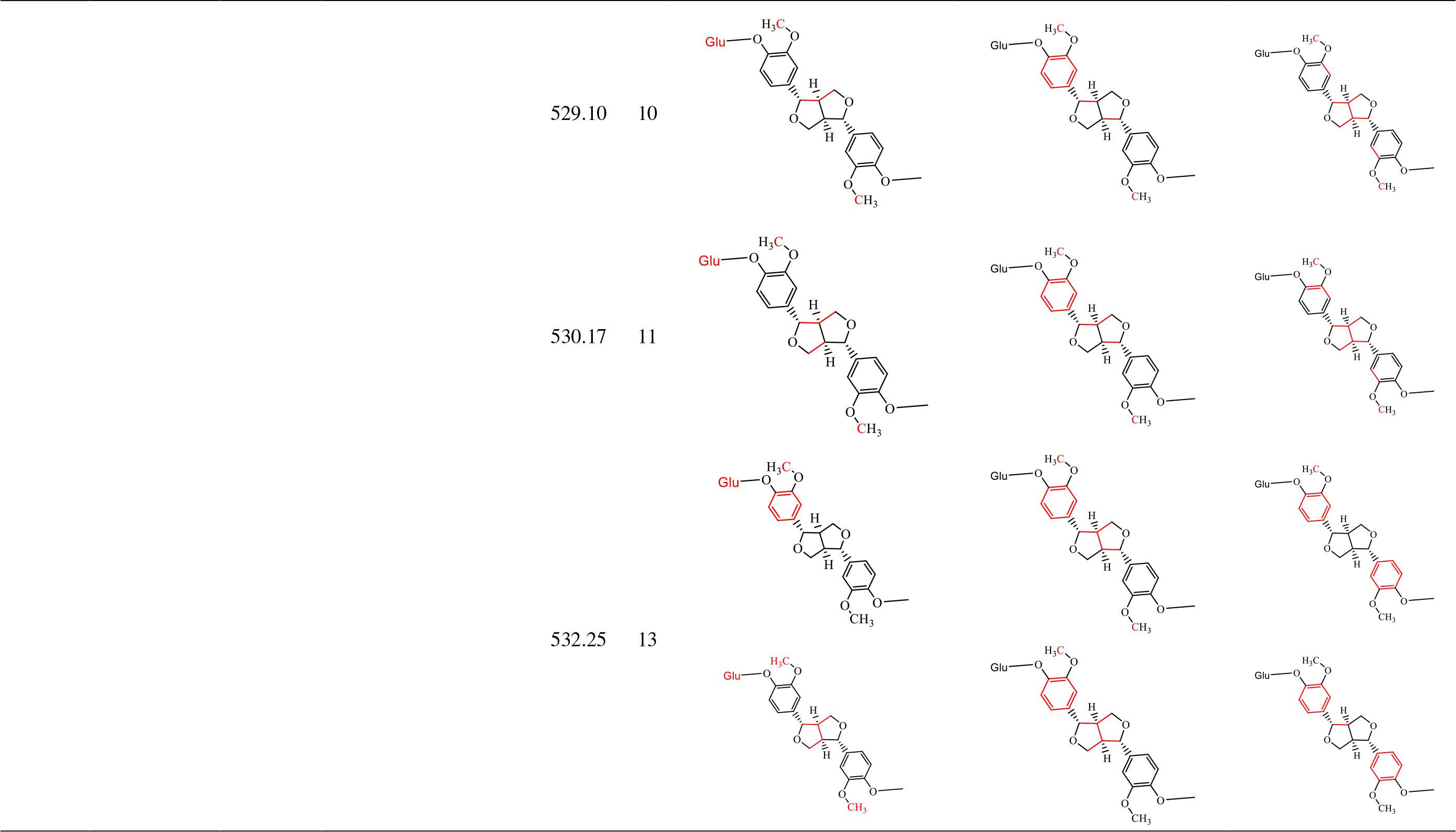

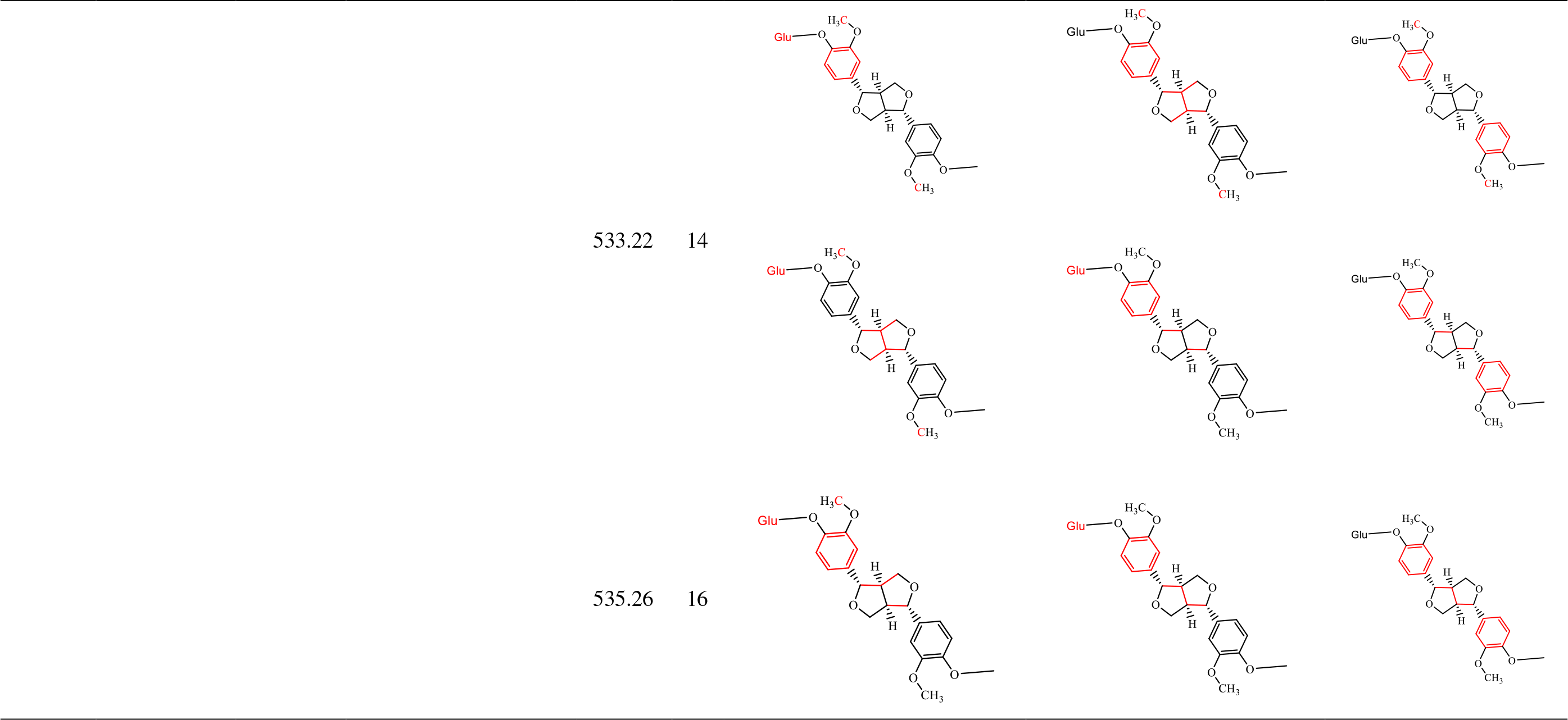

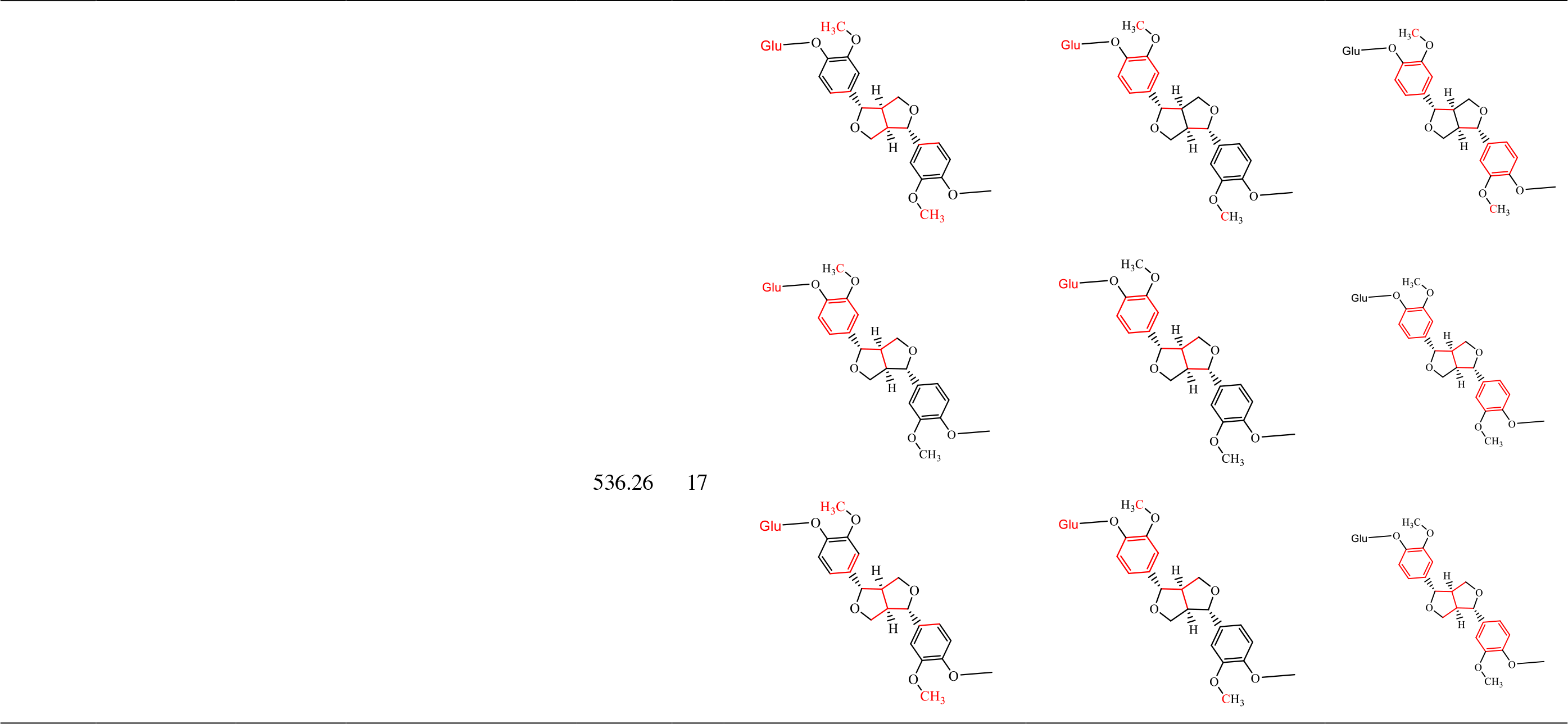

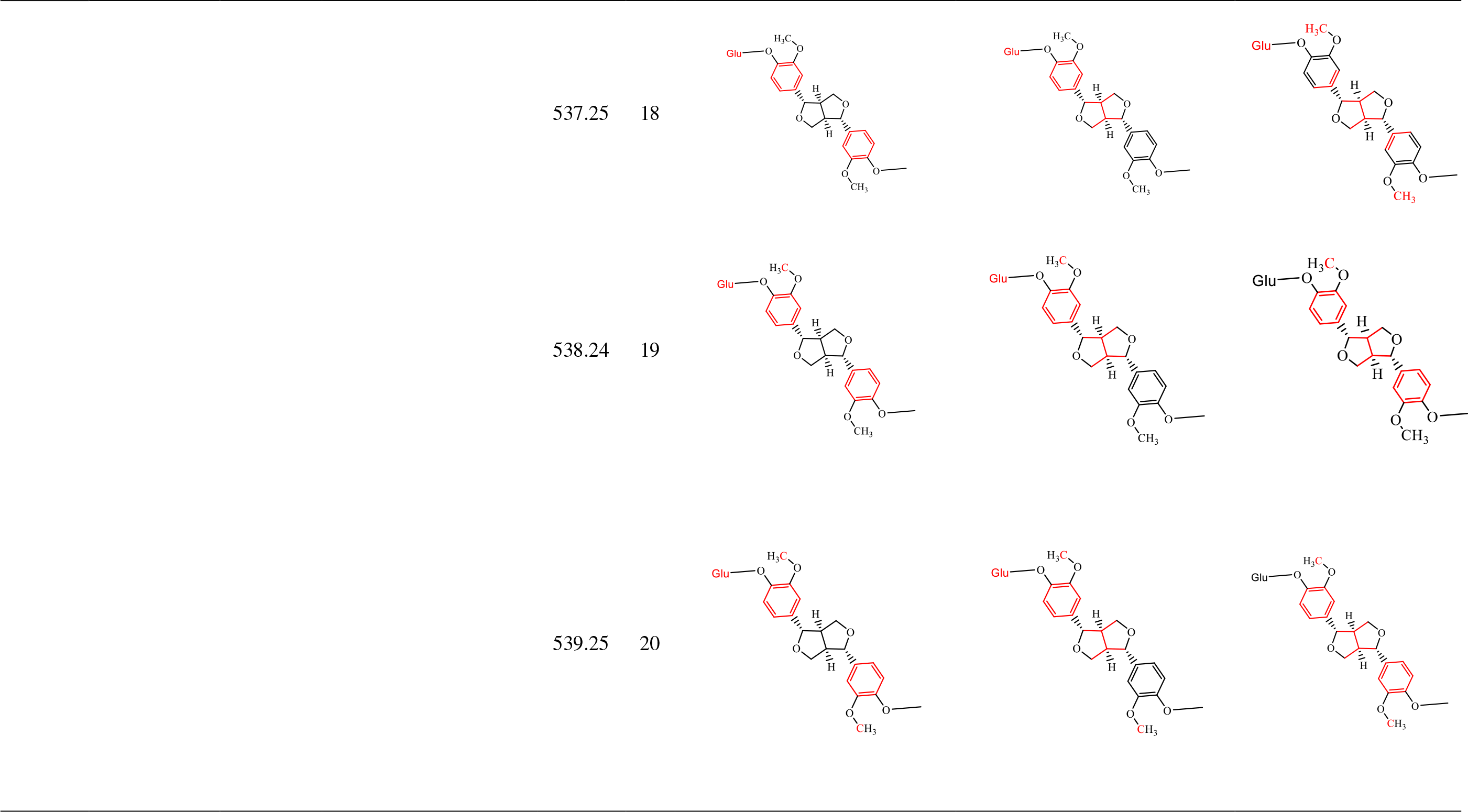

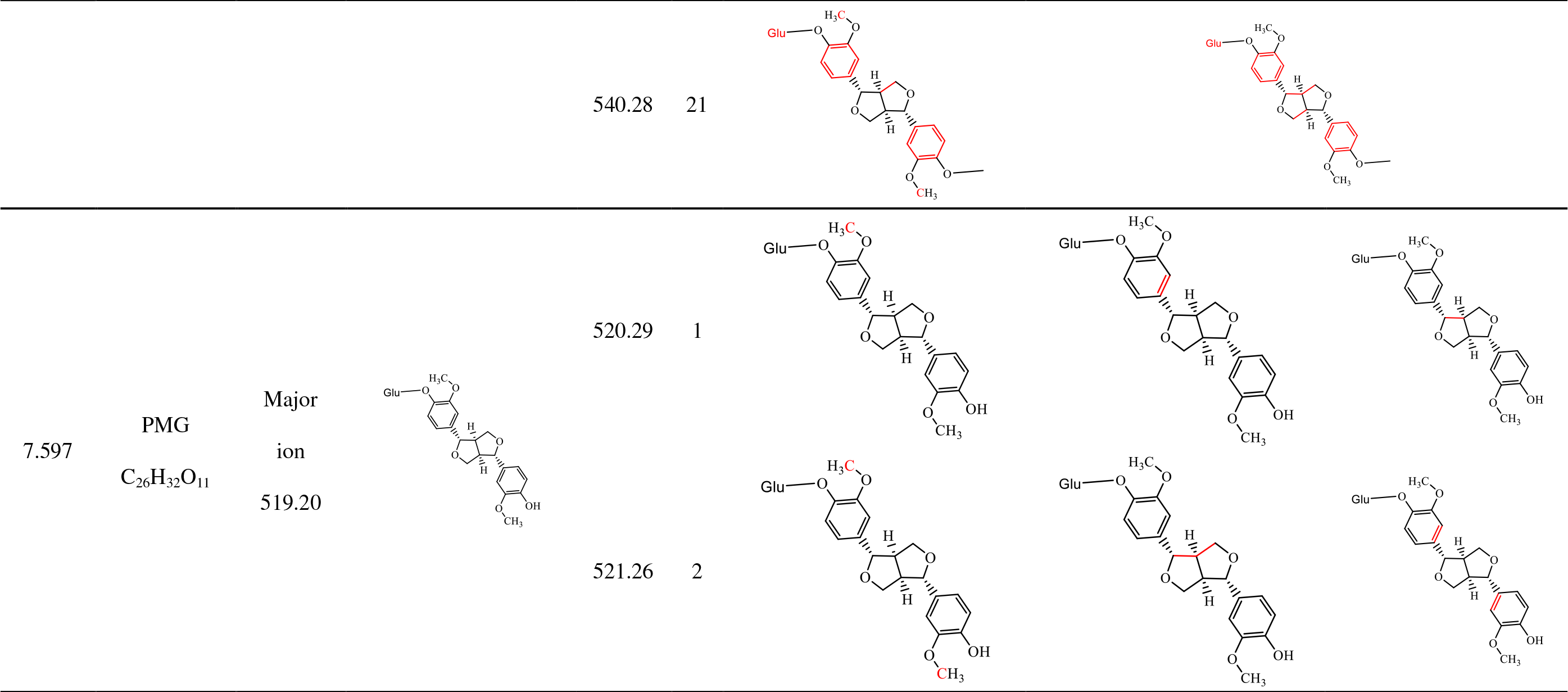

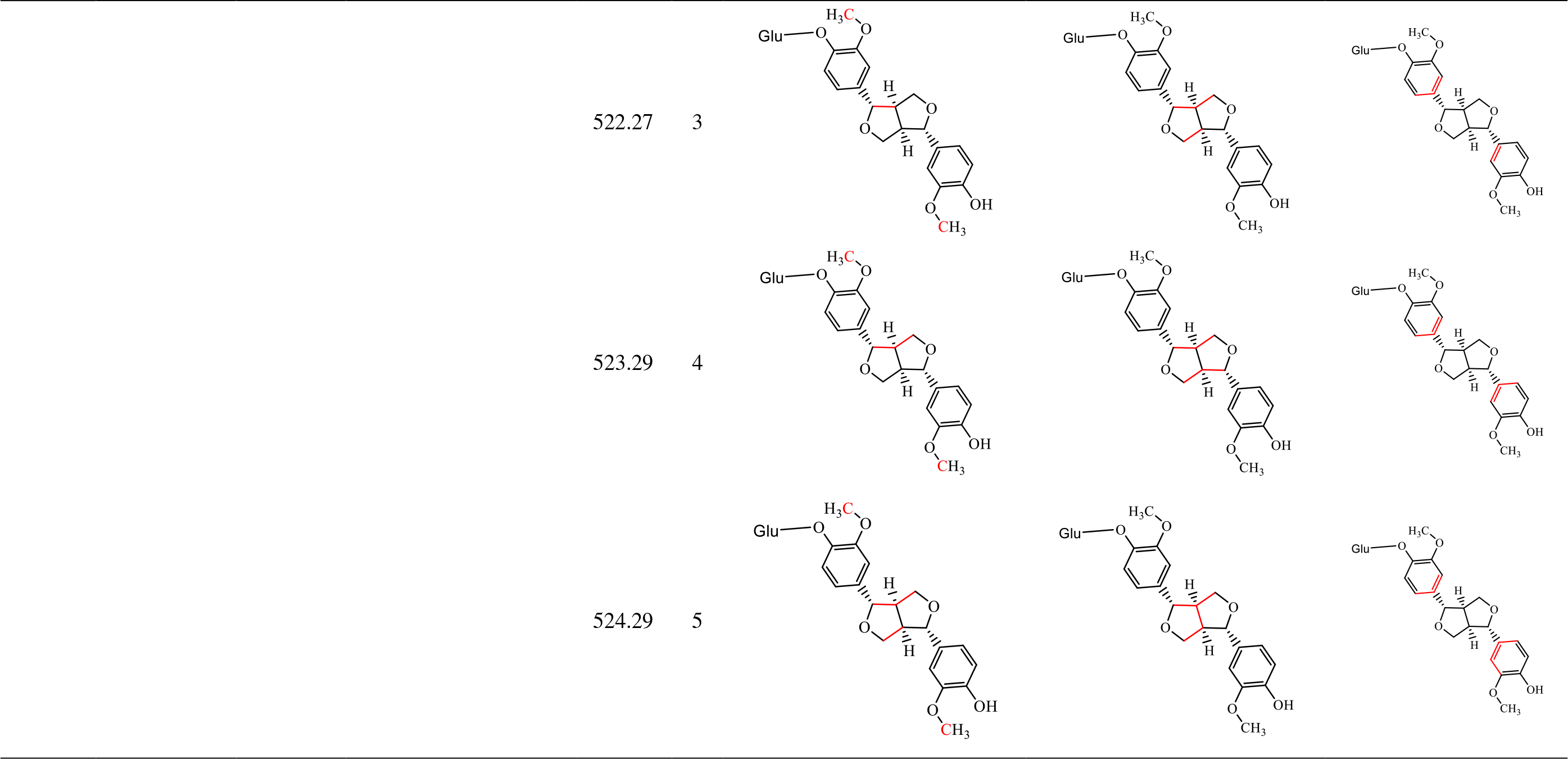

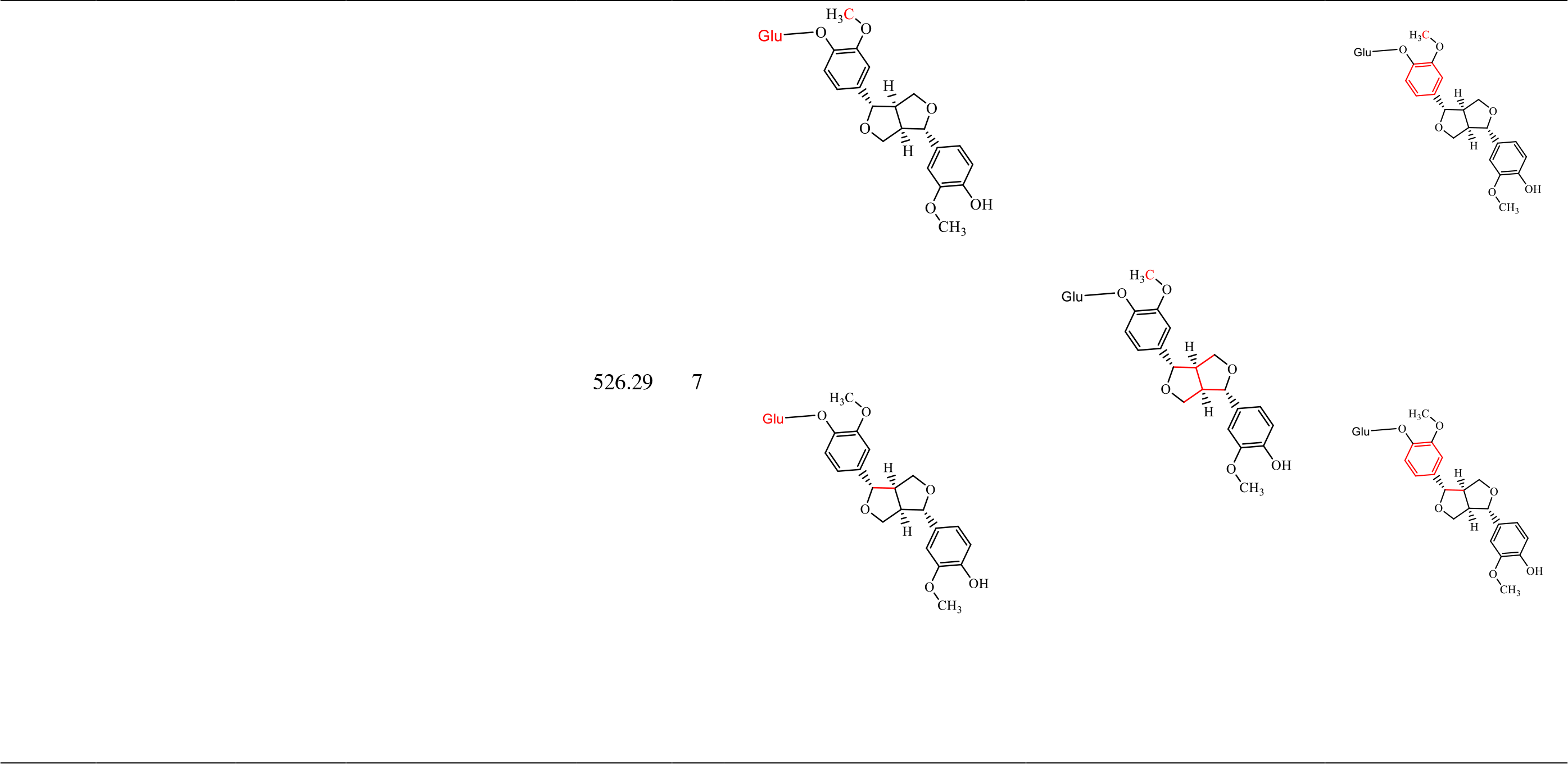

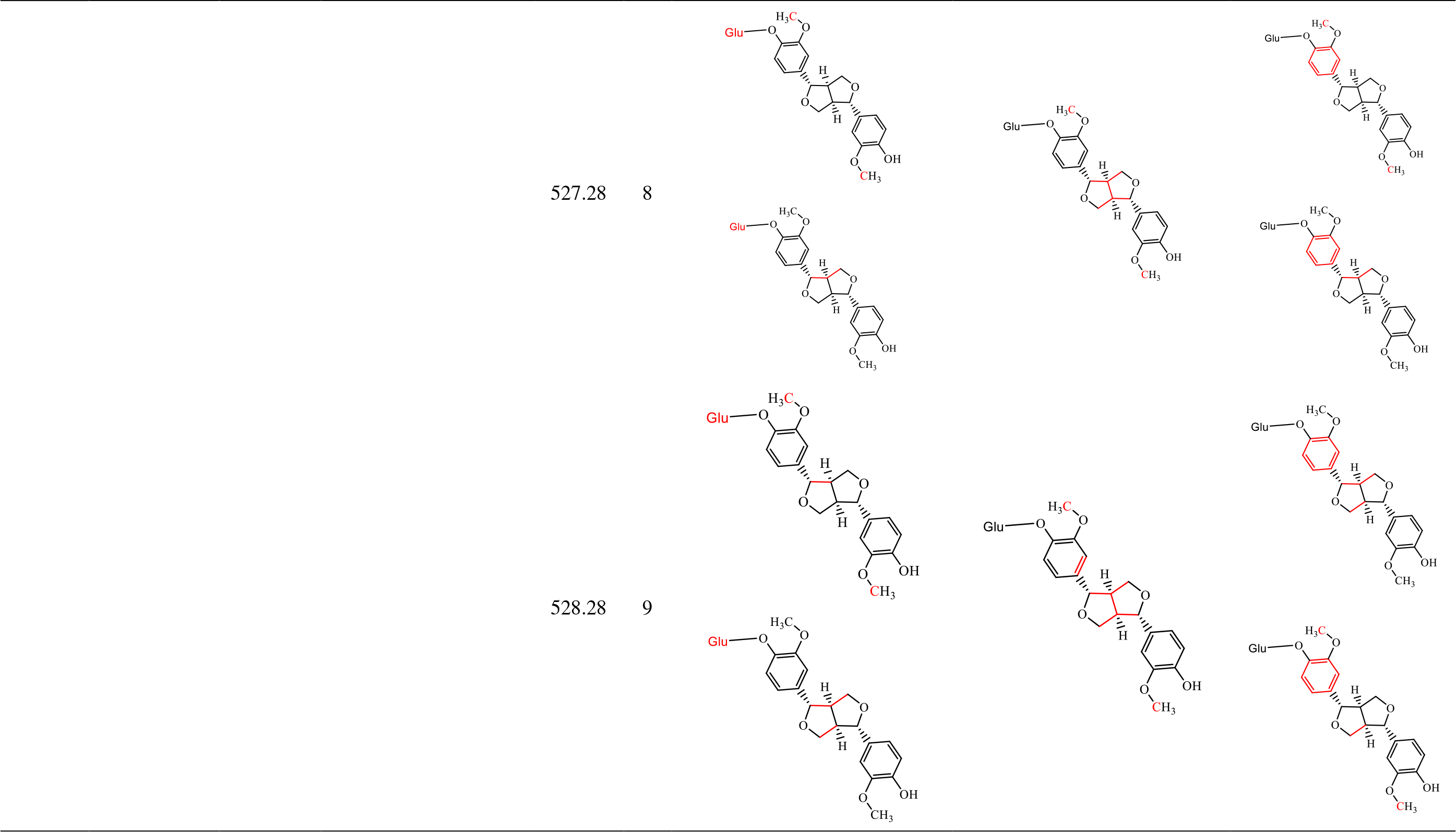

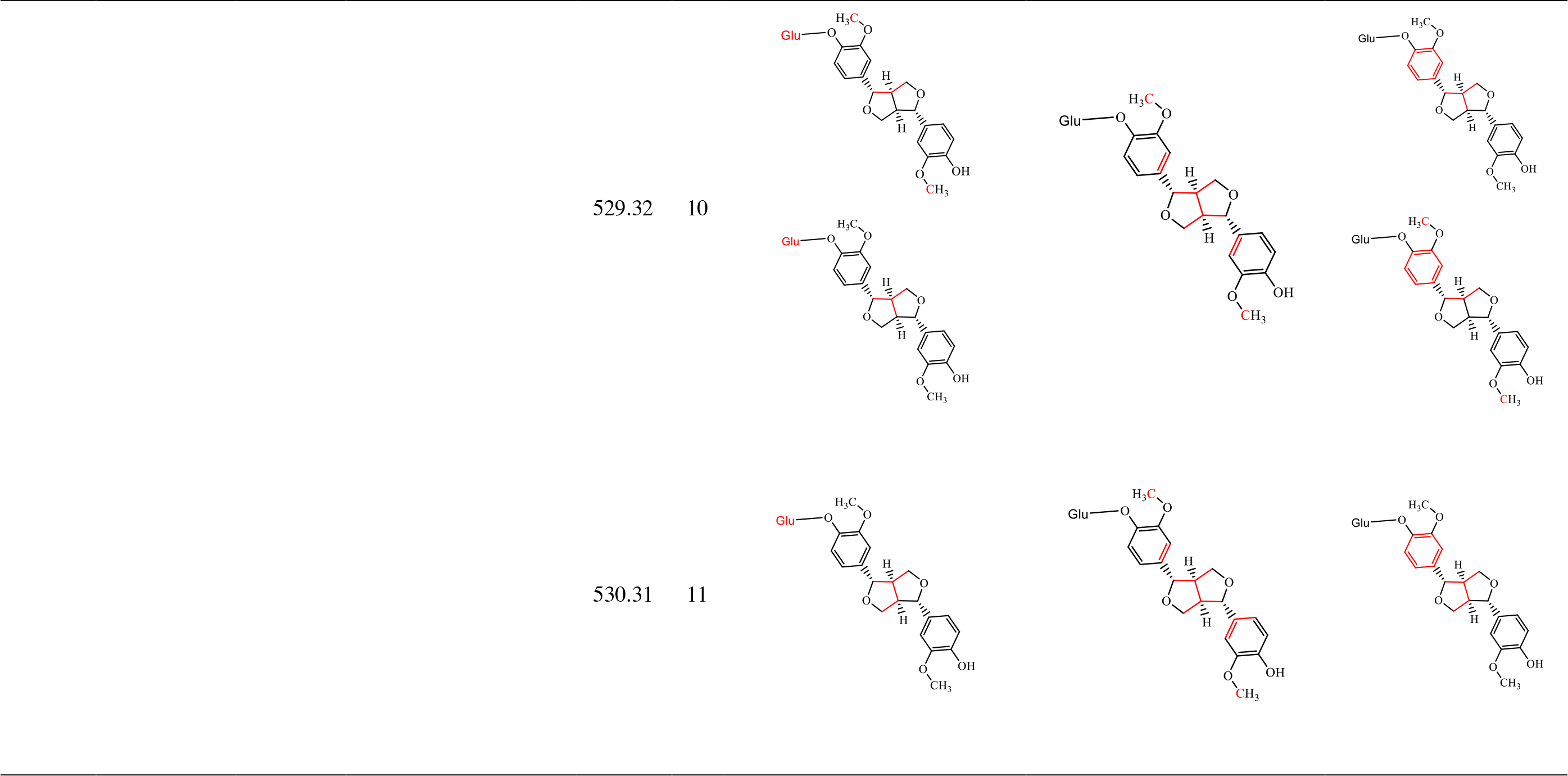

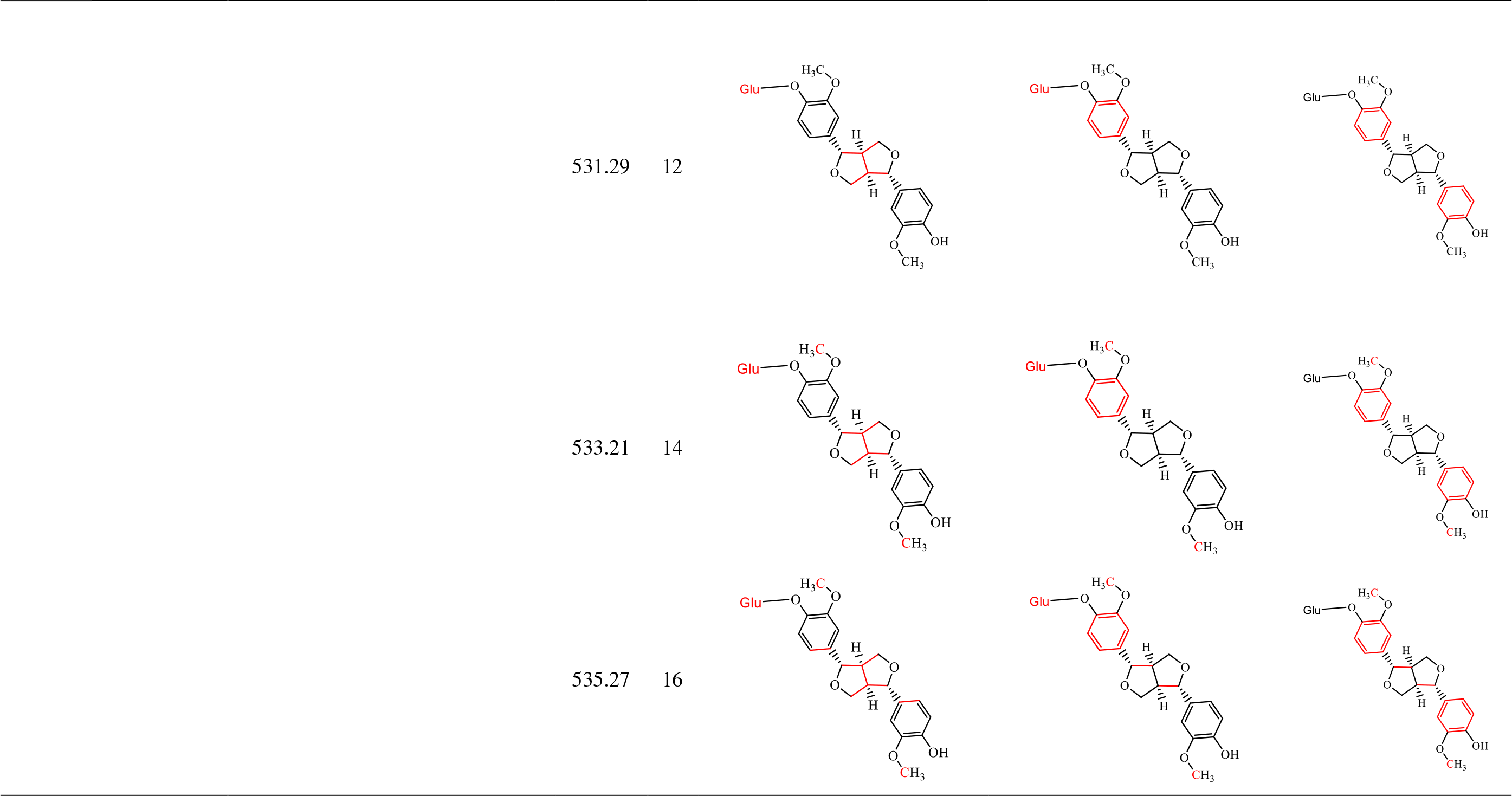

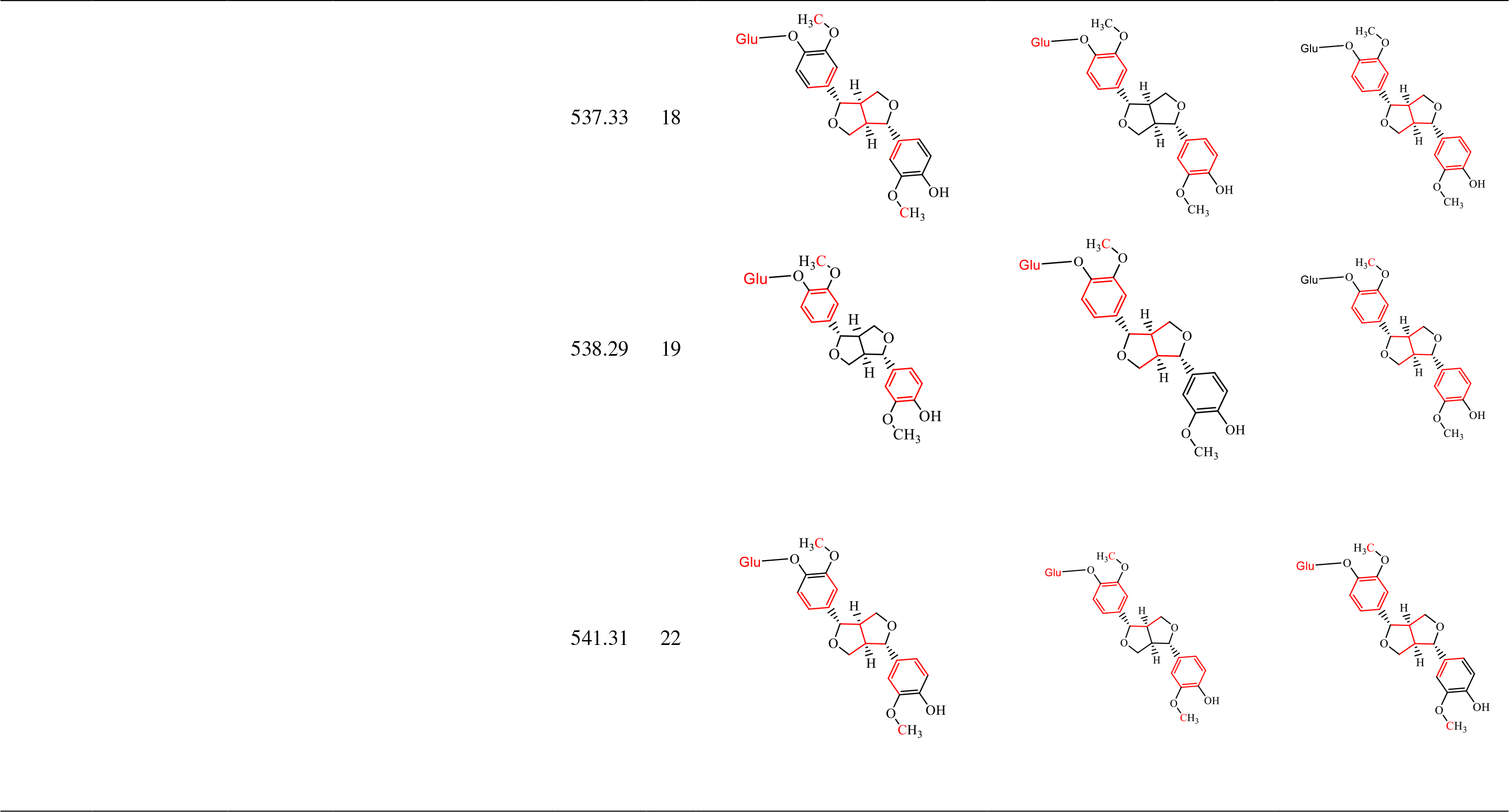

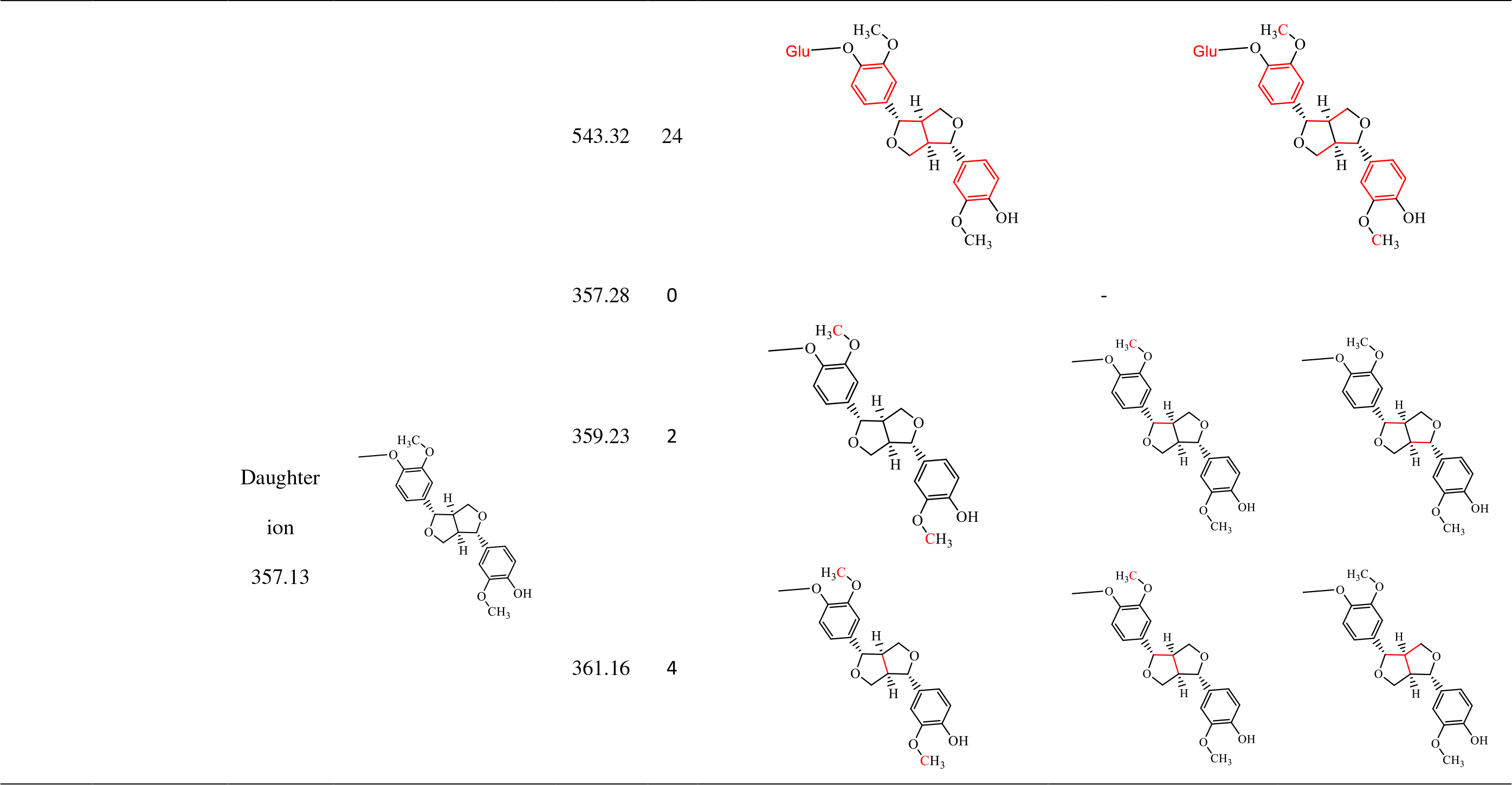

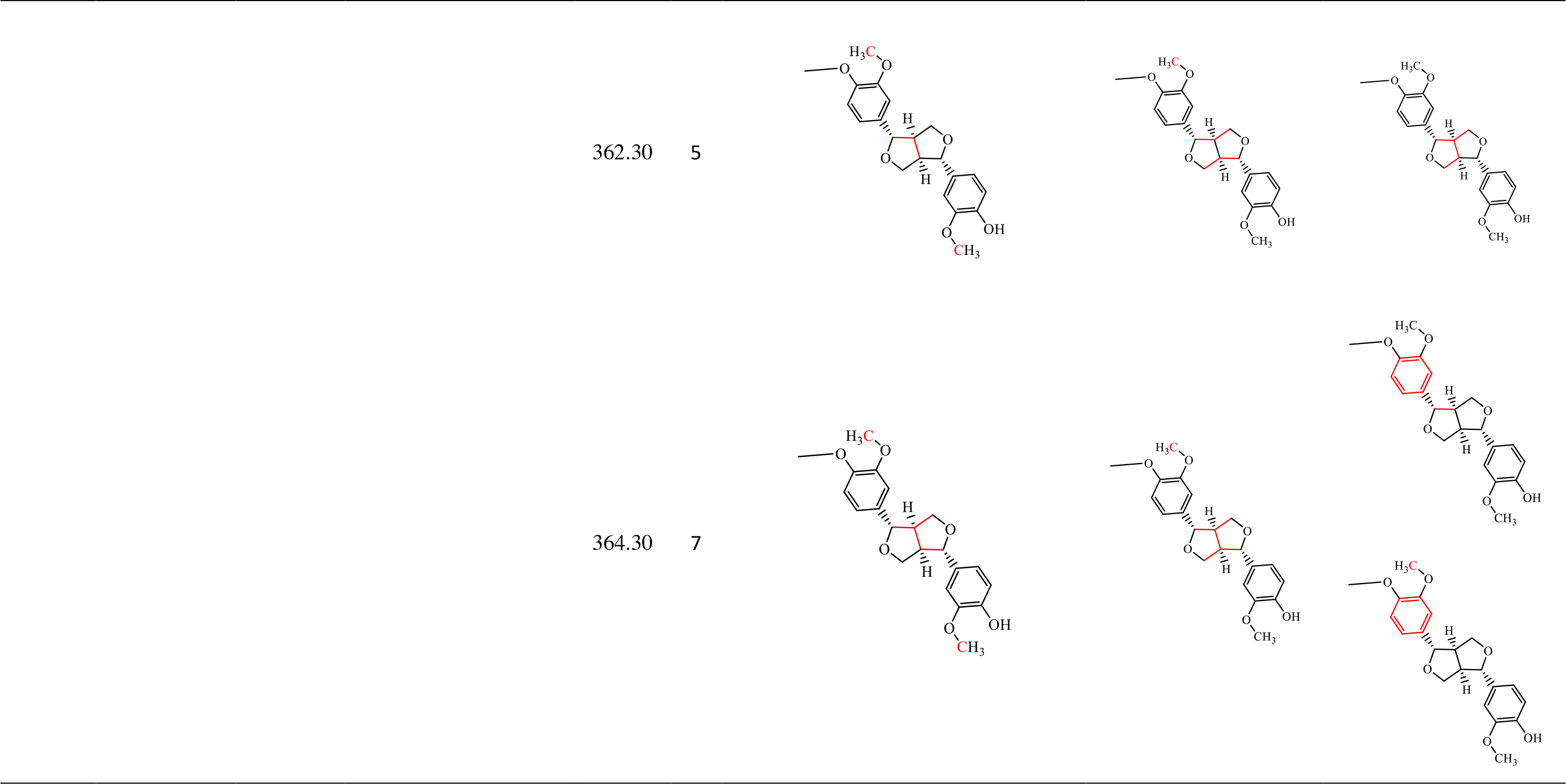

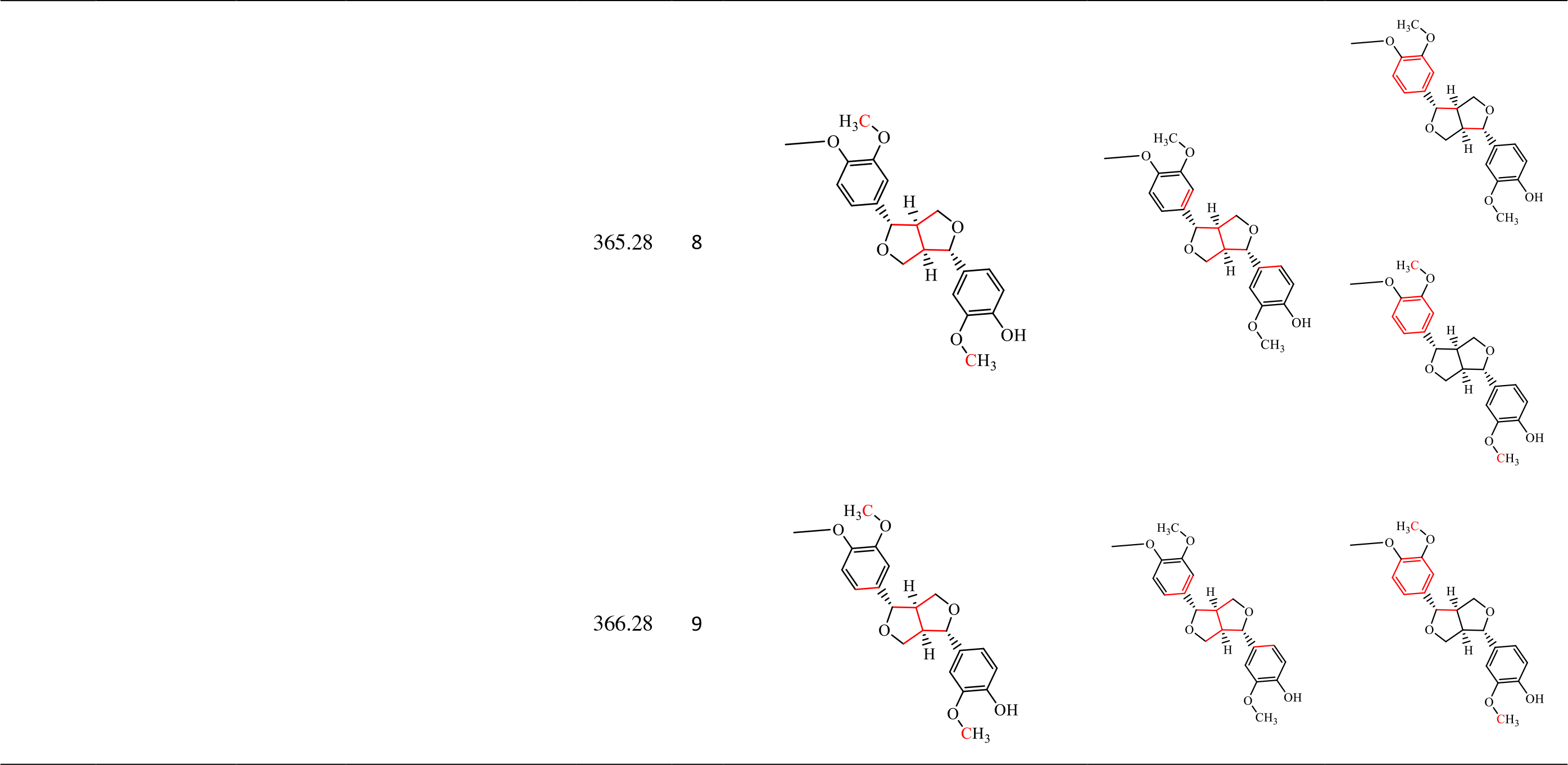

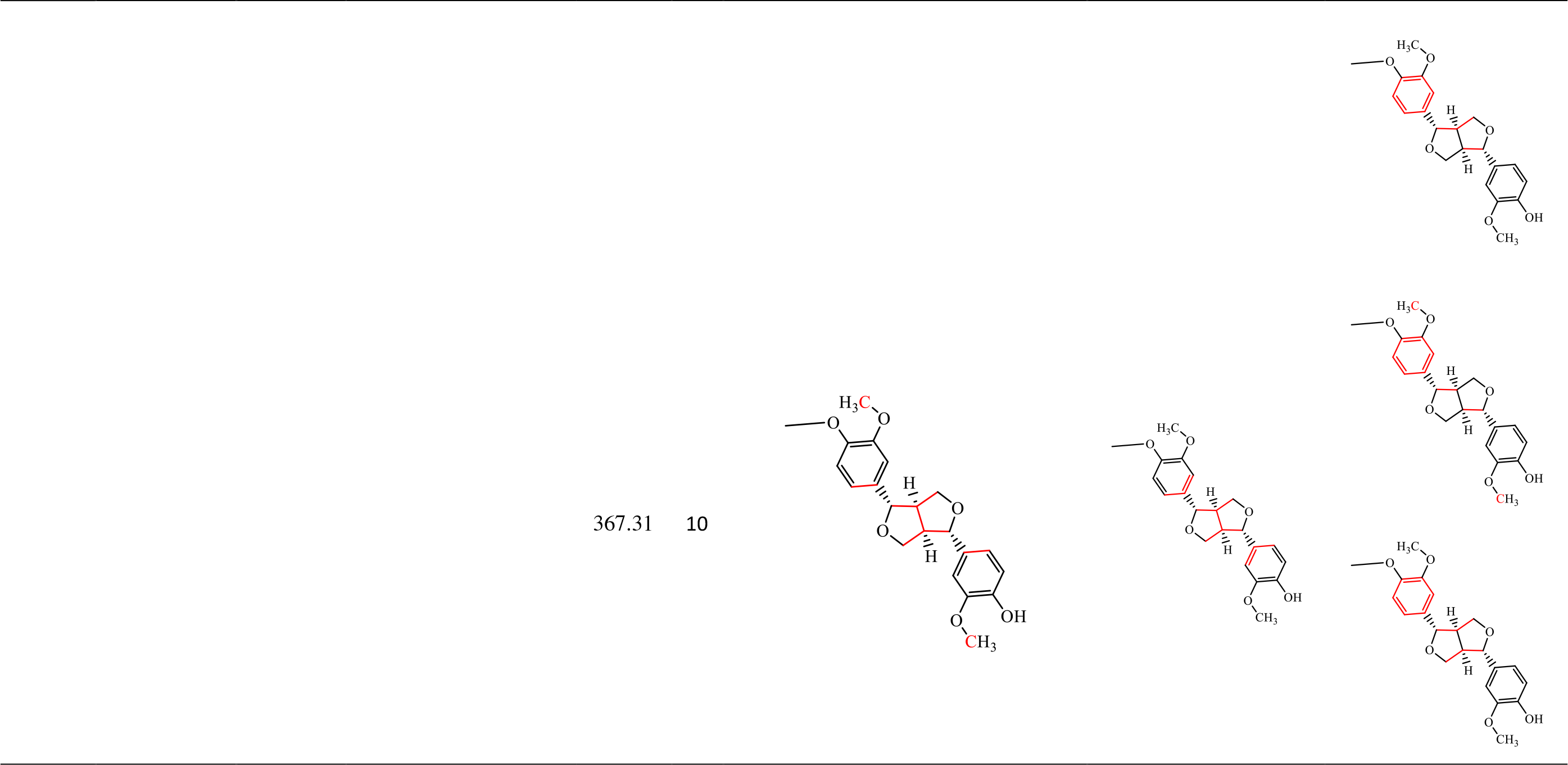

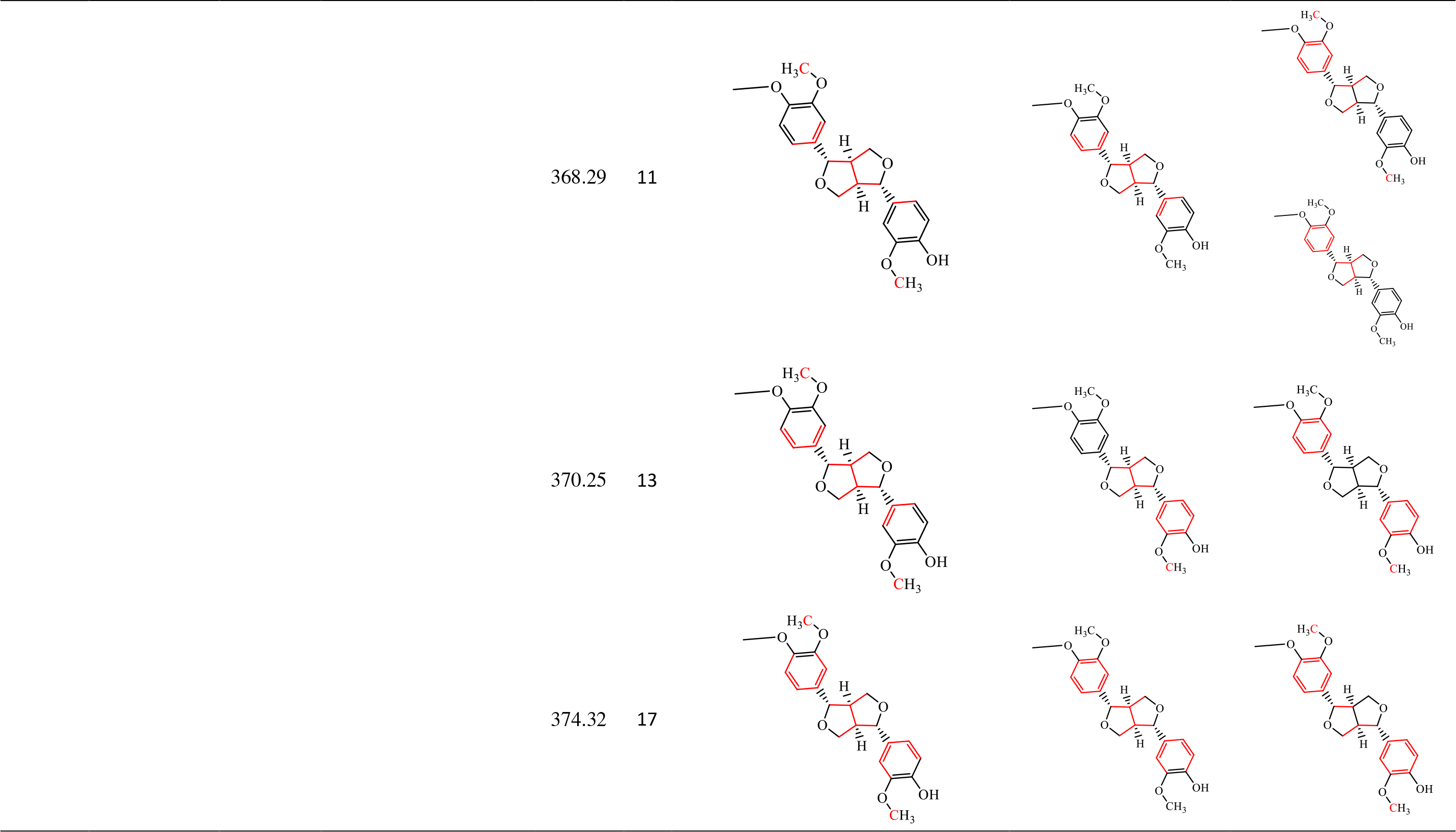

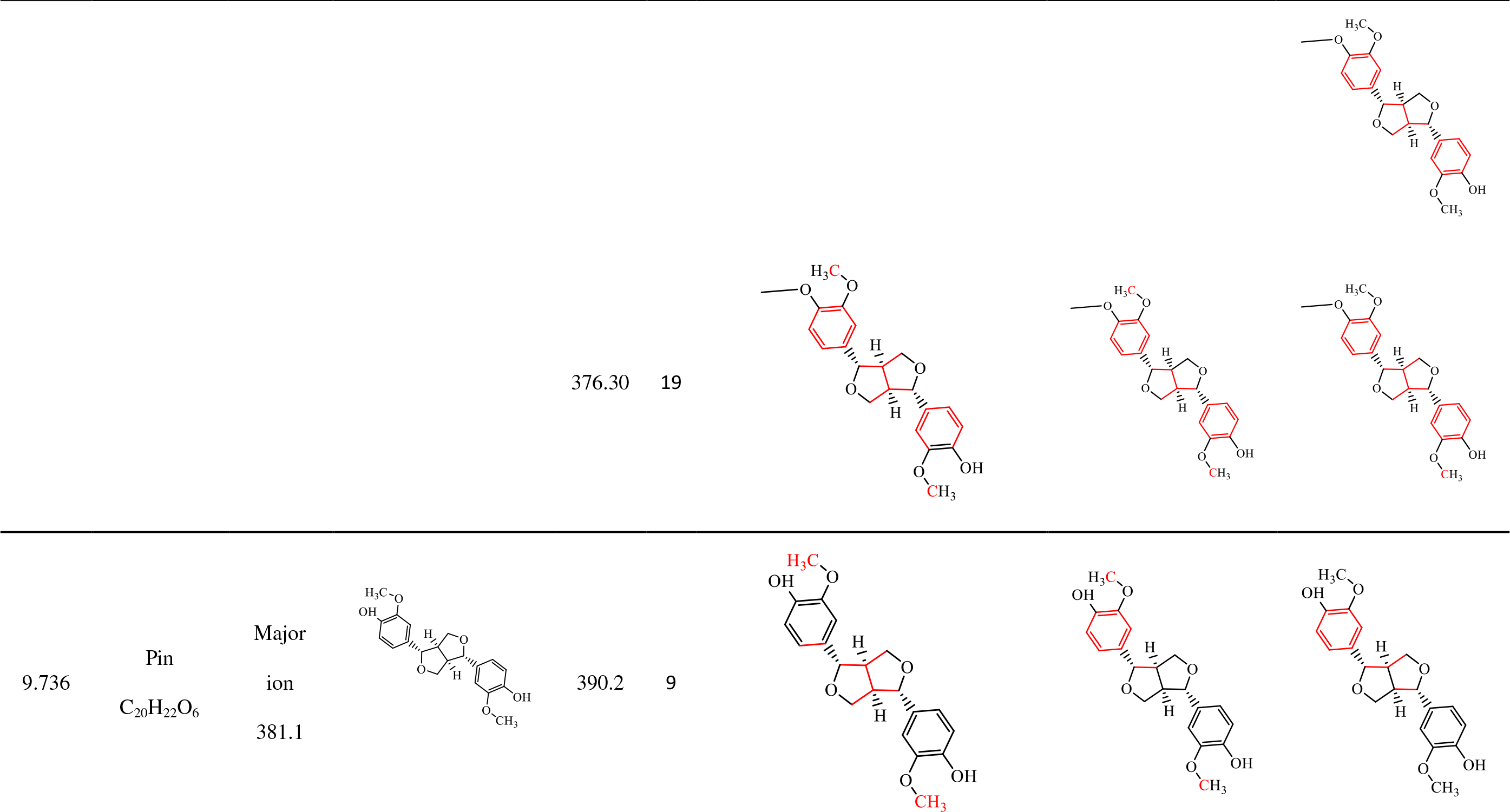

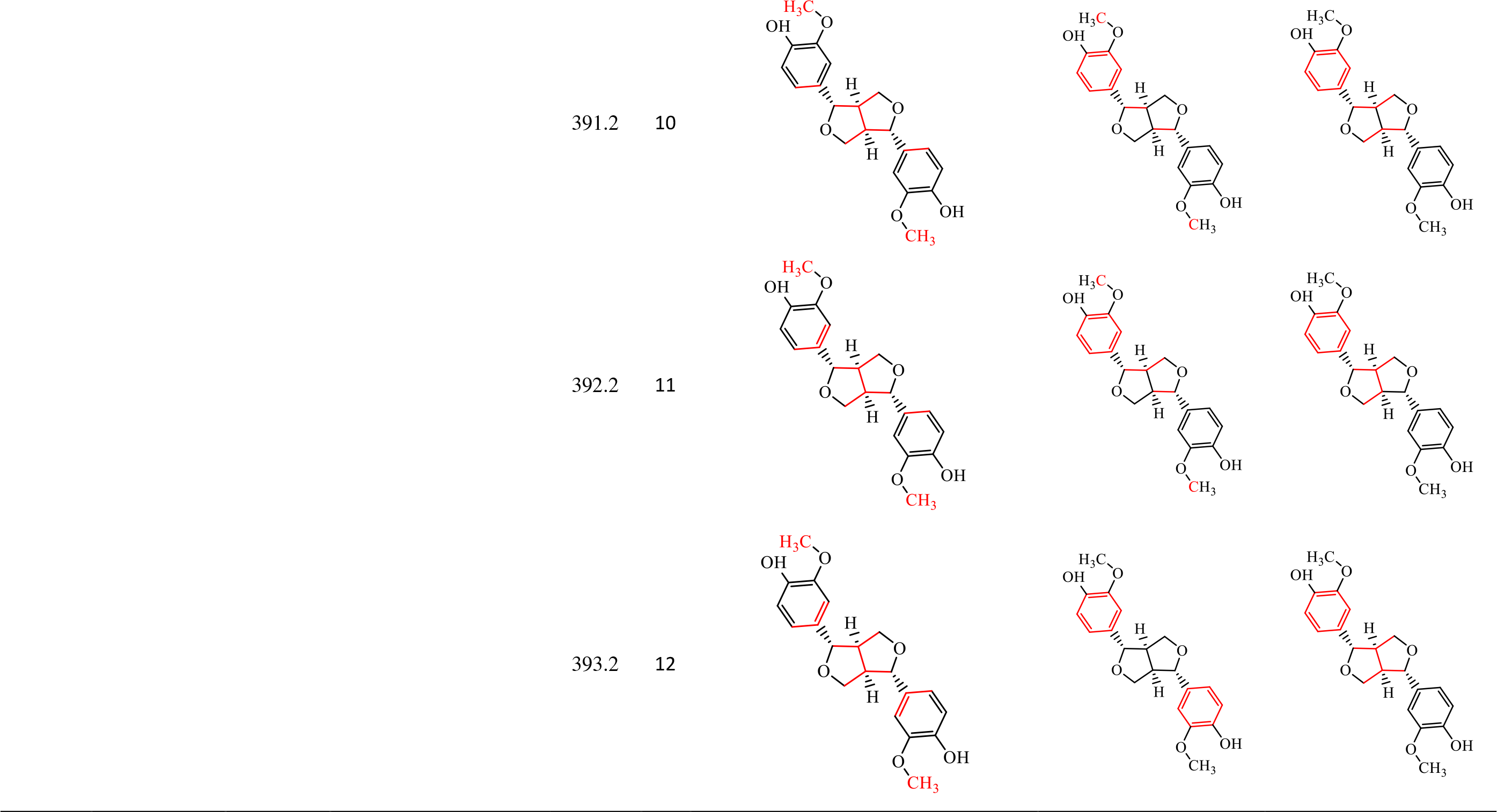

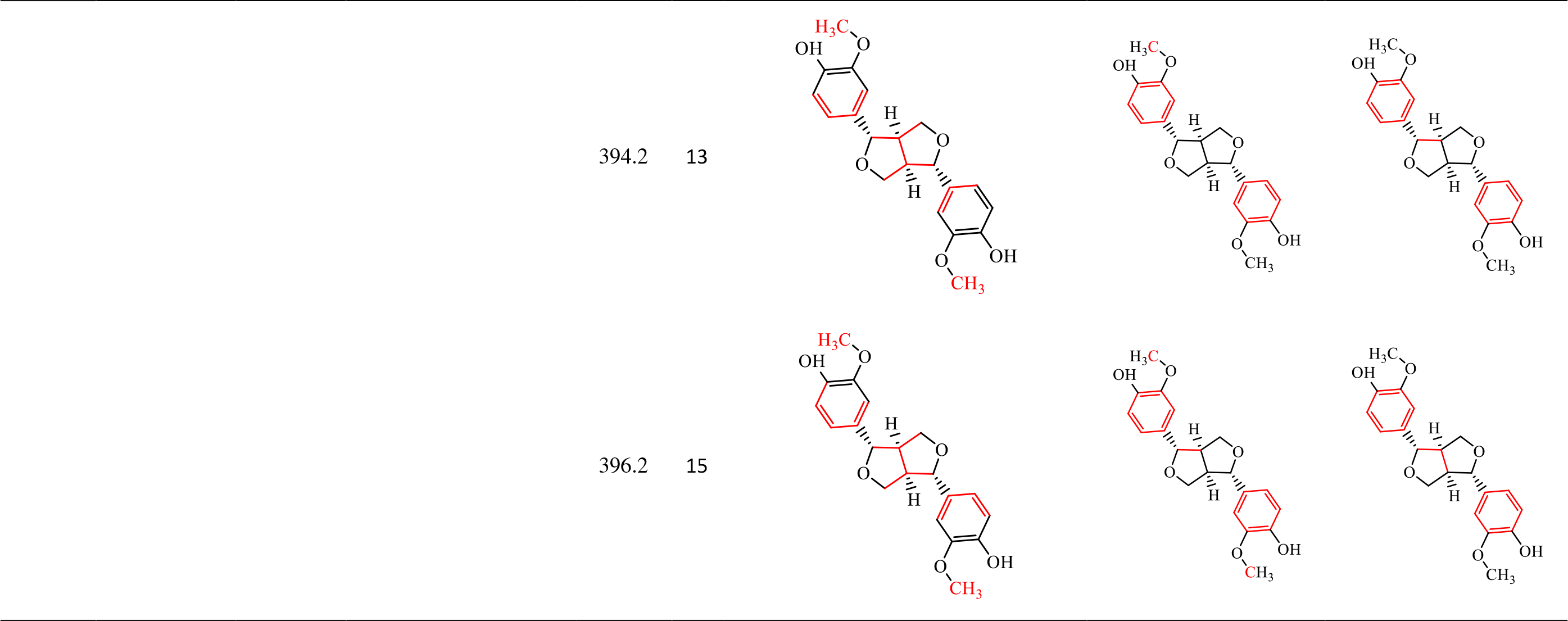

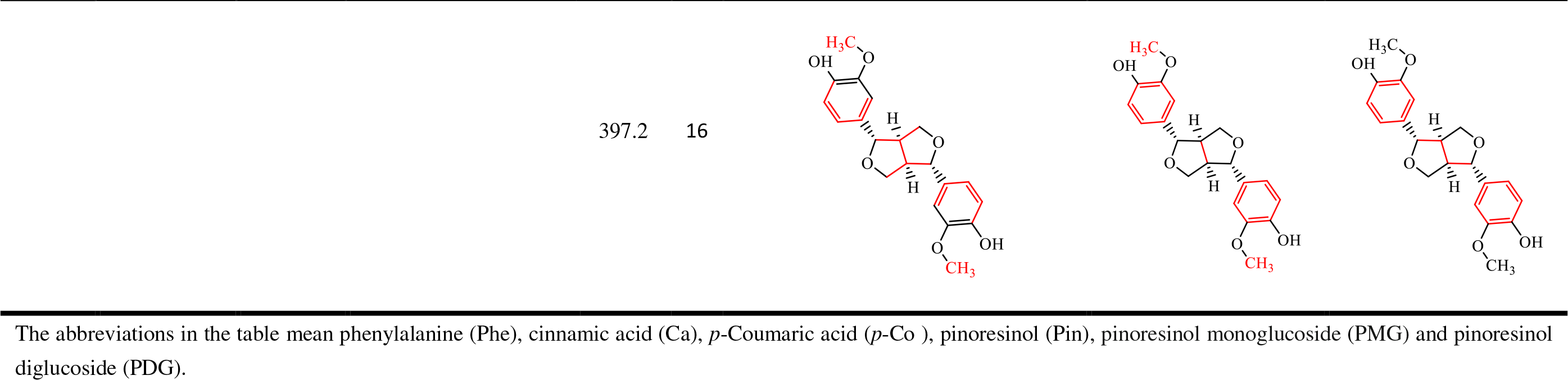

According to the mass spectra of the Phe (C_9_H_11_NO_2_) standard with a RT of 2.06 min, ^13^C labeled Phe was detected in the conversion system with [^13^C_6_]-labeled glucose as the sole substrate at a molecular weight of m/z=168.06, 169.06, 170.07, 171.07, 172.07, and 173.08 (Fig. 9A), indicating four, five, six, seven, eight, and nine ^13^C in ^13^C-labeled Phe, respectively. This result illustrates that there were four, five, six, seven, eight, and nine carbons in the Phe from glucose. The major daughter ions at m/z = 151.04, 107.05, and 108.06 detected in ^13^C-labeled Phe showed that there were four, five, and six ^13^C in Phe, respectively, compared to the daughter ions of the Phe standard (m/z= 147.06, m/z = 103.06). Phe without ^13^C was not detected, indicating that all detected Phe was converted from glucose via the shikimic acid pathway according to KEGG pathway datebase. The possible positions of ^13^C in Phe are summarized in Table 2.

**Fig.9.**
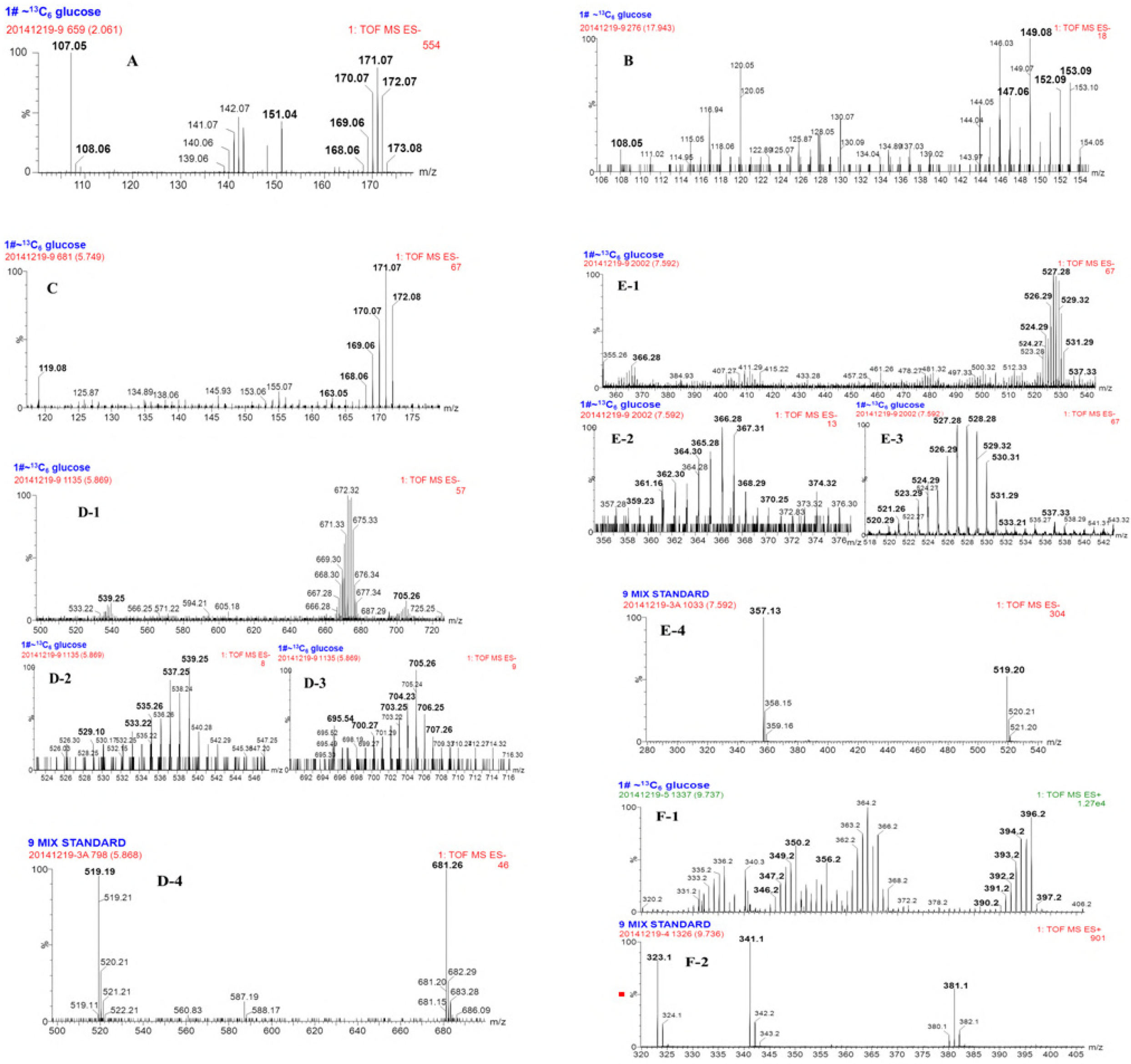
Mass spectrum of phenylalanine, cinnamic acid, *p*-coumaric acid, PDG, PMG, and Pin in the resting cell system using glucose with the ^13^C_6_ stable isotope labeled as the substrate. (A: phenylalanine; B: cinnamic acid; C: *p*-coumaric acid; D: PDG; E: PMG and F: Pin)

Referring to the mass spectra of the Ca standard (C_9_H_8_O_2_, RT= 17.94 min, molecular weight of m/z = 147.05, major daughter ion of m/z = 103.06) (Fig. 3A, B), ^13^C-labeled Ca was detected in the bioconversion system with ^13^C_6_-labeled glucose as the sole substrate (Fig. 9B). ^13^C-Labeled Ca was detected at m/z=147.06, 149.08, 152.09, 153.09, and 154.05, corresponding to zero, two, four, five, and six ^13^Cin the detected molecules, respectively, indicating that they were converted from [^13^C_6_]-labeled glucose. The major daughter ions of ^13^C-labeled Ca were obtained at m/z =108.05, indicating five^13^C in the molecule compared with that of normal Ca (m/z=103.06).Notably, Ca without ^13^C was also detected in the bioconversion system, showing that Ca could also be produced from substrates other than glucose. This was more complex than using Phe as the substrate. The possible positions of ^13^C in Ca are summarized in Table 2.

According to the mass spectra of the *p*-Co standard (C_9_H_8_O_3_, RT= 5.75min, molecular weight of m/z = 163.05, and major daughter ion of m/z = 119.06) (Fig. 4A,B), ^13^C_6_-labeled *p*-Co was detected in the bioconversion system at m/z=163.05, 168.06, 169.06, 170.07, 171.07, and 172.08 (Fig. 9C). Compared to the molecular weight of the *p*-Co standard (m/z= 163.05), the detected ^13^C-labeled *p*-Co indicated that zero, five, six, seven, eight, and nine carbons in *p*-Co came from ^13^C-labeled glucose. The *p*-Co with, five, six, seven, eight, and nine^13^Cmay have been converted from [^13^C_6_]-labeled glucose. The major daughter ion of ^13^C-labeled-Cowas detected at m/z =119.05, which was the same as that of the normal *p*-Co standard (m/z=119.08). Normal *p*-Co was also detected in the bioconversion system, indicating that *p*-Co could also be formed from substrates other than glucose. The possible positions of ^13^C in *p*-Co are summarized in Table 2.

According to the mass spectra of standard PDG (C_32_H_42_O_16_, RT= 5.868 min, molecular weight of m/z = 681.26 and major daughter ion of m/z = 519.19) (Fig. 5A,B), ^13^C-labeled PDG was detected at m/z=695.54, 698.19, 699.27, 700.27, 701.29, 703.25, 704.23, 705.26, 706.25, and 707.26 (Fig. 9D), indicating the occurrence of 14, 17, 18, 19, 20, 21, 22, 23, 24, 25, and 26 ^13^C in PDG, respectively. Glucose may be the sole glycosyl donor in the biosynthesis of PDG by *Phomopsis* sp. XP-8 (Zhang et al. 2015a, b), so there should be more than 12 ^13^C in PDG. As expected, more than 14 ^13^C were detected in ^13^C-labeled PDG. Therefore, it was confirmed that glucose was the sole glycosyl donor for PDG biosynthesis. The maximum number of ^13^C was detected in PDG. If the two glycosides of PDG were all ^13^C, the other 14 ^13^C would from the C-skeleton of the Pin structure; if all C-skeletons of the Pin structure were formed of ^13^C, there would be only one ^13^C-labeled glycoside in PDG. Therefore, glucose not only provided glycoside groups to PDG, but also provided the core Pin structure. The
 possible positions of ^13^C in PDG are summarized in Table 2.

According to the mass spectra of the PMG standard (C_26_H_32_O_11_, RT= 7.597 min, molecular weight of m/z = 519.20,and major daughter ion of m/z = 357.13)(Fig. 6A,B), ^13^C-labeled PMG was detected at a molecular weight of m/z=520.29, 521.26, 522.27, 523.29, 524.29,526.29, 527.28, 528.28, 529.32, 530.31, 531.29, 533.21, 535.27, 537.33, 538.29, 541.31, and 543.32(Fig. 9E), corresponding to 1, 2, 3, 4, 5, 7, 8, 9, 10, 11, 12, 14, 16, 18, 19, 22, and 26^13^C in PMG, respectively, compared with that of PMG (m/z=519.20).One glycosideis present in the structural formula of PMG. If the glycoside came from glucose, there should be at least six^13^Cin PMG. The occurrence of a molecule with less than six ^13^C PMG indicates that the PMG glycoside could have been converted from another substrate, instead of the added [^13^C_6_]-labeled glucose. However, the detection of26^13^C in PMG illustrates that the PMG glucoside could also be converted from [^13^C_6_]-labeled glucose. The major daughter ions of ^13^C-labeled PMG were detected at m/z = 151.04, m/z= 357.28, 359.23, 361.16, 362.30, 364.30, 365.28, 366.28, 367.31, 368.29, 370.25, 374.32, and 376.30 (Fig. 9E-2), indicating 0, 2, 4, 5, 7, 8, 9, 10, 11, 13, 17, and 19^13^C in PMG, respectively, after a comparison to the PMG standard (m/z= 357.13). The major daughter ion of PMG (m/z= 357.13) indicated the molecular weight of the core structure of PMG without the glycoside (20 C). This finding indicates that the core structure of PMG may have partly originated from [^13^C_6_]-labeled glucose. The possible positions of ^13^C in PMG are summarized in Table 2.

According to the mass spectra of the Pin standard (C_20_H_22_O_6_, RT= 9.736 min, molecular weight of m/z = 381.1 and major daughter ion of m/z = 341.1)(Fig. 7A,B), ^13^C-labeled Pin was detected at m/z=390.2, 391.2, 392.2, 393.2, 394.2, 396.2, and 397.2 (Fig. 9F), indicating 9, 10, 11, 12, 13, 15, and 16^13^Cin the detected Pin, respectively, compared with the Pin standard (m/z=381.1). There are 20 C in the molecular formula of Pin (C_20_H_22_O_6_). The maximum of 16^13^Cwasdetected in the formed Pin, indicating the [^13^C_6_]-labeled glucose partly contributed to the formation of Pin. The major daughter ions of ^13^C-labeled Pin were detected at m/z = 346.2, 347.2, 349.2, 350.2, and 356.2 (Fig. 9F-2), indicating 5, 6, 8, 10, and 15 ^13^C in the detected Pin, respectively, compared with the Pin standard (m/z= 341.1). This finding illustrates that the core Pin structure was partly converted from[^13^C_6_]-labeled glucose. The possible positions of ^13^C in Pin are summarized in Table 2.

Taken together, the possible biosynthetic pathways for PDG, PMG, and Pin are summarized in Fig. 10. The mass flow from [^13^C_6_]- Phe to[^13^C_6_]-Ca, [^13^C_6_]-*p*-Co, and[^13^C_12_]-Pin was verified by the experiments using [^13^C_6_]-labeled Phe as the sole substrate. The mass flow from[^13^C_6_]- glucose to[^13^C_6_]-Phe, [^13^C_6_]-Ca, [^13^C_6_]-*p*-Co, [^13^C_12_]-Pin, [^13^C_18_]-PMG, and[^13^C_24_]-PDG was verified by the data obtained using [^13^C_6_]- glucose as the sole substrate (Fig. 10A).

**Fig.10.**
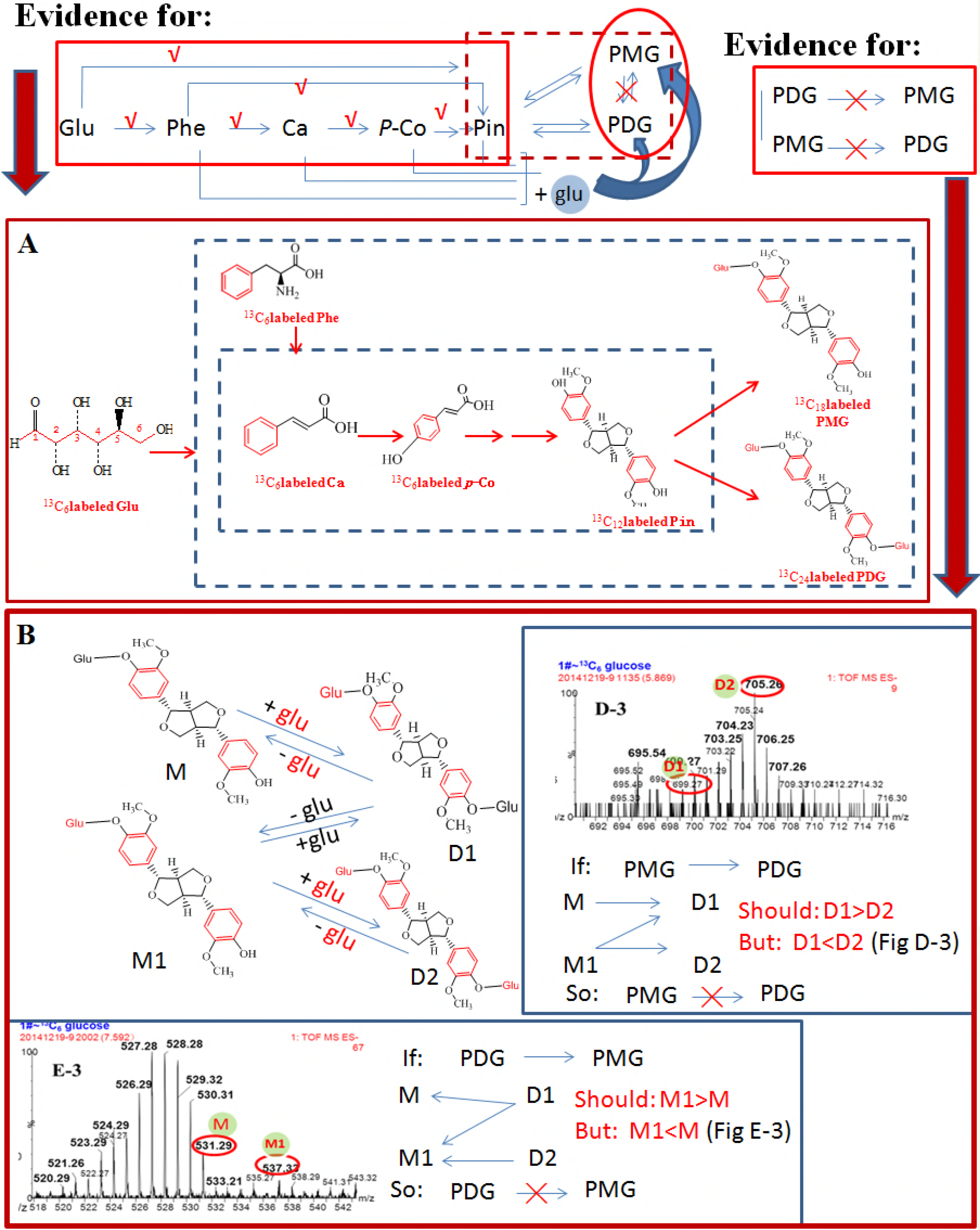
Evidence for a PDG and PMG bioconversion pathway in *Phomopsis* sp. XP-8. The abbreviations in the figure indicate PMG with normal glycoside (M), PMG with^13^C_6_ labeled glycoside (M1), PDG with one ^13^C_6_ labeled glycoside (D1), PDG with two ^13^C_6_ labeled glycoside (D2). ^13^C_6_ labeled glycoside (Red font glu), normal glycoside (Black font glu). √means the pathway was confirmed and ╳means the pathway does not exist in *Phomopsis* sp. XP-8.

### Possible pathways for biosynthesis of PDG and PMG

Two structures of PMG were detected: one was [^13^C_12_]-PMG with two benzene rings converted from ^13^C-labeled glucose and a normal glycoside (M, m/z 531.29), and the other was [^13^C_18_]-PMG with both benzene ring structures converted and a glucoside from ^13^C-labeledglucose (M1, m/z 537.33). Similarly, two PDG structures were detected: one was [^13^C_18_]-PDG with a two benzene ring structure and one glycoside converted from ^13^C-labeledglucose (D1, m/z 699.27); the other one was [^13^C_24_]-PDG with two benzene rings and two glycosides from ^13^C-labeledglucose (D2, m/z 705.26).

If PMG was the direct precursor of PDG, M would be converted to D1 by bonding one [^13^C_6_]-labeled glycoside through glycosylation; M1 could be converted to D1 by bonding one normal glycoside through glycosylation and to D2 by bonding one [^13^C_6_]-labeled glycoside. If this is true, D1 would have two glucoside sources, whereasD2 would have only one glucoside source. Therefore, the concentration of D2 should be lower than D1. However, the data show that the relative abundance of D2 (m/z=705) was much higher than that of D1 (m/z=699). Therefore, PMG was not the precursor of PDG.

In contrast, if PDG was the direct precursor of PMG, D1would be converted to M by hydrolyzation of one [^13^C_6_]-labeled glycoside and to M1 by hydrolyzation of one normal glycoside; D2 would be converted to M1 by hydrolyzation of one [^13^C_6_]-labeled glycoside. If this is true, M1 would have two glycoside sources, where as M would have only one source. The concentration of M should be lower than M1. However, the data show that relative abundance of M (m/z = 531.29) was higher than that of M1 (m/z=699). Therefore, PDG was not the precursor of PMG.

## Discussion

The ^13^C stable isotope labeling method was successfully used in this study to verify the phenylpropanoid-pinoresinol and biosynthetic pathway its glycosides in *Phomopsis* sp. XP-8 during mass flow. It was very important to verify the occurrence of this pathway in microorganisms for the first time. Stable-assisted metabolomics are an efficient way to trace and identify bio-transformed products and the metabolic pathways involved in their formation, such as understanding the fate of organic pollutants in environmental samples (19). This is the first time that this method has been used to verify the occurrence of phenylpropanoids in a microorganism. Compared with previous studies using precursor feeding, detection of enzyme activity, and genomic annotation, this is the first time this pathway has been illustrated by credible visual evidence. More importantly, it is the first time that differences between the PDG and PMG biosynthetic pathways have been verified.

The results obtained in this study verify the existence of the phenylpropanoid-lignan metabolic pathway in *Phomopsis* sp. XP-8. Genomic annotation is an efficient way to discover the pathways that are normally difficult to reveal by metabolic and enzymatic evidence due to low intermediate accumulation, low end-product production, and silent gene expression under normal conditions. This method has been successfully used to identify the existence of a phenylpropanoid metabolic pathway in *Aspergillus oryzae* (28), and the molecular genetics of naringenin biosynthesis, a typical plant secondary metabolite in *Streptomyces clavuligerus*(29), and the occurrence of the phenylpropanoid-lignan pathway in *Phomopsis* sp. XP-8 (17). However, further evidence is still needed to verify gene functions and identify the key metabolites. This study reports the existence of thephenylpropanoid-lignan pathway *Phomopsis* sp. XP-8during mass flow and identified the metabolites. Further studies are still needed to verify the gene functions.

Additional studies should illustrate the origin of the genes in the phenylpropanoid-lignan pathway of *Phomopsis* sp. XP-8. Horizontal gene transfer (HGT) has long been recognized as an important force in the evolution of organisms (30). HGT occurs among different bacteria and plays important roles in the adaptation of microorganisms to different hosts or environmental conditions (31). More and more evidence for gene transfer between distantly related eukaryotic groups has been presented (30).Therefore, we cannot exclude the possibility that XP-8 may have acquired the genes related to the lignan biosynthetic pathway from its host plant by HGT during long-term symbiosis and evolution. However, further evidence is still needed to verify this proposed process.

The results obtained in this study provide useful information on the biosynthesis of lignans and their glycosides via microbial fermentation. Biosynthesis of lignans is of great interest to organic
 chemists as it provides a model for biomimetic chemistry and has extensive applications (32).Improvement shave been made in the techniques to biosynthesize lignan products by regulating the lignan biosynthetic pathway in trees through genetic modifications (33).However, the lignan biosynthetic pathway has rarely been reported. More importantly, the bioconversion sequence from Pin to PDG and the direct precursor of PDG have remained unclear until now. In previous studies on *Phomopsis* sp. XP-8, the highest production of PDG and PMG did not occur simultaneously (14) and PMG was not the precursor of PDG because PDG production decreased and/or disappeared when PMG yield increased (15). The present study demonstrated that PMG was not the precursor of PDG, and PDG was not the precursor of PMG, indicating that Pin might be converted to PMG and PDG via two different pathways in *Phomopsis* sp. XP-8, which has not been revealed in plants.

Furthermore, this study revealed that the bioconversion of Pin, PMG, and PDG from glucose occurred simultaneously as that from Phe. We found that the benzene ring structure of Phe did not open throughout the entire Pin bioconversion process in *Phomopsis* sp. XP-8 when Phe was used as the sole substrate, indicating that the Pin benzene ring originated from Phe. Glucose was converted to Phe and was the sole glycoside donor for PDG biosynthesis. Therefore, glucose not only participated in the formation of glycosides in PDG, but also provided the PDG benzene ring structure. This is different from that found in plants, indicating there might be some other different pathways to produce these products in *Phomopsis* sp. XP-8.

Not all intermediates in the KEGG-identified plant-lignan biosynthetic pathway related to Pin, PMG, and PDG formation were found in *Phomopsis* sp. XP-8, such as caffeic acid, ferulic acid, and coniferyl alcohol (Fig. 1). This may be because the pathways after *p*-Co are different in XP-8 from those in plants, or the accumulation of these intermediates was too less to be detected. Further studies are needed to verify this hypothesis.

## Conclusion

In summary, the mass flow of the Pin, PMG, and PDG biosynthetic pathway in *Phomopsis* sp. XP-8was verified as the following: starting from[^13^C_6_]- Phe to[^13^C_6_]-Ca, [^13^C_6_]-*p*-Co, and[^13^C_12_]-Pin when there was only Phe as the sole substrate; starting from [^13^C_6_]- glucose to[^13^C_6_]-Phe, [^13^C_6_]-Ca, [^13^C_6_]-*p*-Co, [^13^C_12_]-Pin, [^13^C_18_]-PMG, and[^13^C_24_]-PDG when there was high level of glucose (15 g/L) as the sole substrate (Fig. 10A).

## Acknowledgements

We acknowledge funding by the National Natural Science Foundation of China (grant no. 31471718), the Modern Agricultural Industry Technology System (CARS-30), the National Key Technology R&D Program (2015BAD16B02), the National Natural Science Foundation of China (grant no. 31760446), and the Start-up funding of Shihezi University (RCSX201713), and Key research and development plan of Shaanxi Province (2017ZDXL-NY-0304).

